# Multi-Cellular Network Model Predicts Alterations in Glomerular Endothelial Structure in Diabetic Kidney Disease

**DOI:** 10.1101/2024.12.30.630833

**Authors:** Krutika Patidar, Ashlee N. Ford Versypt

**Affiliations:** Department of Chemical and Biological Engineering, University at Buffalo, The State University of New York, Buffalo, NY 14260, USA; Department of Biomedical Engineering, University at Buffalo, The State University of New York, Buffalo, NY 14260, USA; Institute for Artificial Intelligence and Data Science, University at Buffalo, The State University of New York, Buffalo, NY 14260, USA; Department of Pharmaceutical Sciences, University at Buffalo, The State University of New York, Buffalo, NY, 14215, USA

## Abstract

Diabetic kidney disease (DKD) progression is often marked by early glomerular endothelial cell (GEC) dysfunction, including alterations in the fenestration size and number linked to impaired glomerular filtration. However, the cellular mechanisms regulating GEC fenestrations remain poorly understood due to limitations in existing *in vitro* models, challenges in imaging small fenestrations *in vivo*, and inconsistencies between *in vitro* and *in vivo* findings. This study used a logic-based protein-protein interaction network model with normalized Hill functions for dynamics to explore how glucose-mediated signaling dysregulation impacts fenestration dynamics in GECs. We identified key drivers of fenestration loss and size changes by incorporating signaling pathways related to actin remodeling, myosin light chain kinase, Rho-associated kinase, calcium, and VEGF and its receptor. The model predicted how hyperglycemia in diabetic mice leads to significant fenestration loss and increased size of fenestrations. We found that glycemic control in the pre-DKD stage mitigated signaling dysregulation but was less effective as DKD developed and progressed. The model suggested alternative disease intervention strategies to maintain fenestration structure integrity, such as targeting Rho-associated kinase, VEGF-A, NFκB, and actin stress fibers.

## Introduction

The kidney is a highly vascularized organ that includes different populations of endothelial cells (ECs) with specialized structures and functions [1]. ECs in the kidney microvasculature regulate blood flow, coagulation, inflammation, and vascular permeability [1, 2]. In the functional unit of the kidney, glomerular endothelial cells (GECs) are highly specialized ECs that contribute to the structural and functional integrity of the glomerular filtration barrier in each nephron and support other glomerular cells, such as podocytes and mesangial cells, and the glomerular basement membrane [3, 4].

The structure of GECs consists of transcellular holes, known as fenestrations, and a rich surface of glycocalyx that together contribute to the size- and charge-selective properties of the filtration barrier [1, 5–8]. The endothelial glycocalyx lines the luminal side of the GECs and is also present within the fenestrations [7, 9]. The functional significance of fenestrations is to provide selective passage of proteins, fluid, and small solutes across the GEC barrier without the need for endocytosis or receptor-mediated mechanisms [7, 10]. The fenestrated endothelium regulates the glomerular filtration rate and permeability [5, 9, 10]. Mature GEC fenestrations are generally open and non-diaphragmed, located in the peripheral cytoplasm, and arranged in the non-raft sieve plates [5, 9, 11, 12]. Fenestrations are supported by a fenestrae-associated cytoskeletal ring (FACR) [13, 14]. Fenestrations are structurally supported by a cytoskeletal lattice, wherein structural proteins like spectrin and filamin crosslink with the actin cytoskeleton, which contributes to fenestration formation, membrane integrity, cell-cell interactions, and shape changes [2, 13–15]. Although their protein composition remains mostly unknown, studies have been done to determine the structural composition of fenestrations [13, 16, 17]. Besides structural proteins, vascular endothelial growth factor (VEGF), endothelin-1, and tumor necrosis factor (TNF)-α are among other agents that modulate endothelial fenestration structure and vascular permeability [2, 12]. Shear stress also regulates the production of vasoactive mediators, such as nitric oxide (NO) and endothelin-1, and regulates fenestration structure and vascular tone [18].

GECs are susceptible to injury and dysfunction in kidney diseases. Alterations in the size and density of GEC fenestrations are associated with the disruption in glomerular filtration and progression of diabetic kidney disease (DKD) [9, 10, 19]. DKD is a microvascular dysfunction in the kidneys and is reported in 20–50 % of diabetic patients [20]. DKD is the leading cause of end-stage renal failure and is associated with significantly increased comorbidities and mortality [10, 20]. GEC activation and dysfunction are considered early signs of DKD development and progression [6, 21].

A few recent studies have focused on understanding dysfunction and injury in endothelial cells in a diseased state. Accumulated evidence associates pathways and signaling motifs with actin cytoskeletal rearrangement and, ultimately, morphological changes in fenestrations in LSECs, which are structurally similar to GECs. VEGF, NO, and calcium are linked with the regulation of myosin light chain kinases (MLCK), Rho, and Rho-associated protein kinase (ROCK) [2, 13, 14, 22–24]. Calcium-dependent inactivation of MLCK and Rho/ROCK activation reduces endothelial porosity in LSECs and, in some cases, increases the diameter of fenestrations [13, 14, 25]. Although most previous studies were performed on fenestrated LSECs, some recent studies focused on fenestrated GECs [2, 10]. Despite some progress toward a mechanistic understanding of markers and pathways associated with GEC dysfunction, research gaps remain in understanding the relation between signaling molecules and mechanical cues in GECs that cause structural deformation. Using the information about pathways in other fenestrated ECs is a promising strategy for understanding these pathways’ potential impacts on the structural or functional stability of GECs before they can be validated experimentally in GECs.

Computational network models have proven useful in mechanistically linking signaling cues to cellular dysfunction and activation. Previously, we [26] and others [27–29] used computational modeling to demonstrate the intracellular signaling and intercellular crosstalk among pro-inflammatory mediators, pro-angiogenic factors, immune cells, and endothelial cells. We previously developed a logic-based ordinary differential equations (LBODEs) model to predict the effects of high glucose and inflammation on macrophage and GEC activation and signaling dysregulation observed *in vitro* associated with early-stage DKD [26]. Others have also effectively used complex network models to study different macrophage phenotypes in response to mixed pro- and anti-inflammatory stimuli [29, 30]. Several large-scale logic-gated network models have effectively determined signaling components and network topology that regulate cell phenotype, function, and structure in other tissues [31, 32].

In this study, we used an LBODES model of protein-protein interaction to study the development and progression of DKD. Here, we modified and extended our previously developed LBODEs model [26] to include essential proteins and interactions that potentially modulate fenestration density and size in GECs *in vivo*. We re-fitted and validated the influential parameters in the extended LBODEs model using published experimental data of observed changes in fenestration size and number in diabetic mice. Using this extended LBODEs model, we analyzed network motifs and dynamics under varying glucose stimuli and perturbed protein activity. We have identified potential strategies for therapeutic interventions to reduce fenestration loss across early to advanced stages of DKD.

## Methods

### Network assembly

We modify our previously developed protein signaling network (PSN) between macrophages and GECs [26] to include relevant pathways and proteins associated with changes in GEC fenestrations (Fig 1). Several experimentally determined effects are considered to extend the previous PSN [26] to the network in Fig 1. The new regulatory nodes and interactions in Fig 1 are derived from proteins and mechanisms proposed in published experiments and hypotheses with fenestrated LSECs [13, 14, 16, 24, 33] and GECs [2]. The new regulatory nodes include IL-1R, RhoA-associated kinase (Rho/ROCK), myosin light chain (MLC), MLC phosphatase (MLCP), MLC kinase (MLCK), phosphorylated myosin light chain (pMLC), stressed actin fibers, relaxed actin fibers, fenestration number, and fenestration diameter. Abbreviations are defined in Table A1 in S1 Appendix.

**Fig 1.**
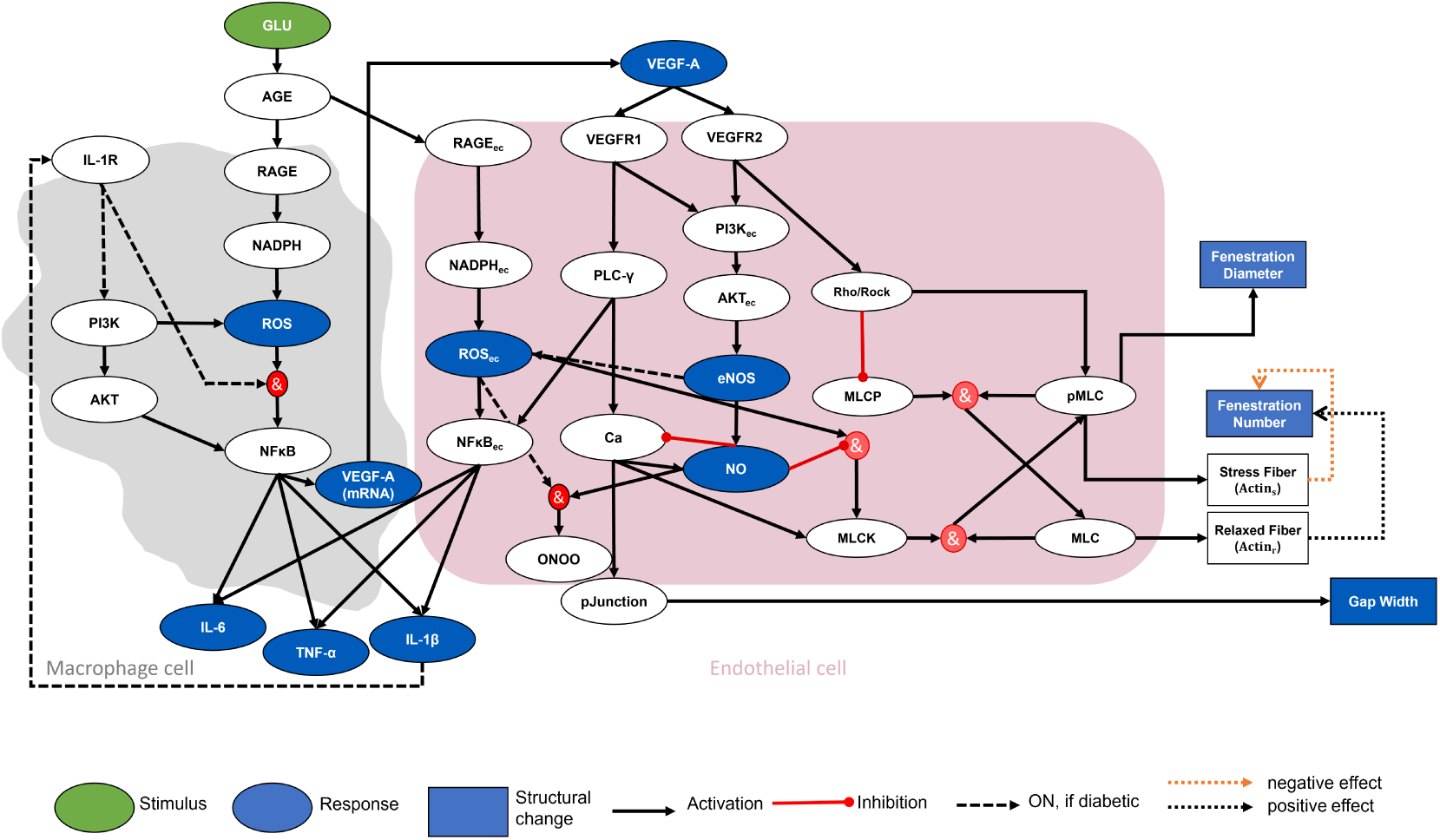
Multi-cellular protein interaction network of *in vivo* crosstalk between macrophages (gray area) and glomerular endothelial cells (pink area) is stimulated with glucose. The green oval is the input node, the blue ovals are the output nodes, and the white ovals are the regulatory nodes. Structural changes in glomerular endothelial cells are shown as blue squares. The black solid arrows are activating interactions, the red edges with dots at one end are inhibiting interactions, and the gray dashed arrows are interactions active for diabetic subjects. Open dotted arrows in orange and black represent negative and positive effects on fenestration size, respectively. Red circles indicate logic *AND* gates. An *OR* logic rule connects two or more edges to a subsequent node throughout the network unless indicated otherwise by an *AND* logic gate. The subscript “ec” denotes an intracellular species expressed in endothelial cells. IL-6, TNF-α, IL-1β, and VEGF-A are protein levels expressed in extracellular space. ROS, ROSec, VEGF-A (mRNA), and NO are expressed within the cells. The Gap Width node denotes a fractional change in the intercellular gap between GECs. The pJunction node represents the phosphorylated junction protein levels. Stress fibers and relaxed fibers represent different forms of actin fibers. AGE: advanced glycation end products. AKT: serine/threonine-specific protein kinases. Ca: calcium. eNOS: endothelial nitric oxide synthase. IL: interleukin. MLC: myosin light chain. MLCK: myosin light chain kinase. MLCP: myosin light chain phosphatase. NADPH: nicotinamide adenine dinucleotide phosphate. NFκB: nuclear factor kappa B. NO: nitric oxide. ONOO: peroxynitrite. PI3K: phosphoinositide 3-kinases. PLC-γ: phospholipase C gamma. pMLC: phosphorylated myosin light chain. RAGE: receptor of advanced glycation end product. ROCK: RhoA-associated kinase. ROS: reactive oxygen species. TLR: toll-like receptor. TNF-α: tumor necrosis factor-alpha. VEGF: vascular endothelial growth factor. VEGFR: vascular endothelial growth factor receptor.

Unlike the previous PSN, which was stimulated by a fixed glucose and lipopolysaccharide stimulus [26], the extended model is stimulated by a dynamic glucose concentration in diabetic mice and an endogenous inflammatory stimulus (IL-1β) indirectly regulated by glucose. Glucose is the only independent stimulus in the extended network model (Fig 1) and is responsible for initiating a phenotypic switch in macrophages, activating GECs, and initiating downstream signaling dysregulation. IL-1β activates its receptor IL-1R on macrophages in the extended PSN.

VEGF, a pro-angiogenic factor, increases EC porosity and permeability in different cell types [2, 14, 34–37] and promotes maintenance of fenestrations via NO-dependent or NO-independent pathways. We link VEGF-A, VEGF receptor, NO, ROSec, and calcium in the previous PSN with additional nodes in the network. VEGF receptor 2 (VEGFR2) mediates Rho/ROCK activation, leading to defenestration in LSECs [2, 13, 36]. Treatment with reactive oxygen species or nitrogen species increases fenestration diameter and decreases fenestration number [14]. The calcium level, regulated by calcium membrane channels and pumps, causes a cascade of cellular mechanisms that drive local changes in the cytoskeleton and result in actin contraction [14, 38]. The exact mechanism of action of NO on fenestration has not been shown. However, it has been demonstrated that eNOS-derived NO shows a positive effect on LSEC fenestration maintenance [36]. It was proposed that activation of the NO-dependent cGMP pathway reduces the activation of MLCK [14]. The local balances regulating the calcium, ROS, NO, and VEGF levels in different parts of the cell control the dynamics of fenestrated LSEC [14], which is included in the extended network. The activity of MLCK is increased by calcium and protein kinase C (PKC)-mediated phosphorylation [14]. The link between MLCK and calcium is considered in the extended network. Moreover, indirect inhibition of MLCK through a calcium-dependent or calcium-independent manner reduced endothelial porosity and increased fenestration diameter in a few cases [25]. Indirect inhibition of MLCK via NO and ROS activation is included in the extended network. MLCP maintains the balance of phosphorylation or dephosphorylation of MLC. MLCK and MLCP, together, keep the balance between pMLC and MLC protein levels (Fig 1). The Rho/ROCK pathway activates pMLC protein and inhibits MLCP protein [14, 25]. pMLC protein increases contractile forces in the actin cytoskeleton structure, which we model as activation of stress fibers in the extended network (Fig 1). Other agents such as PKA, PKG, and PKC—not explicitly included in the extended network—may also activate pMLC; however, they are not as potent as the Rho/ROCK pathway [14]. RhoA regulates the assembly of contractile actin bundles and actin filaments, and negative regulators of Rho/ROCK have resulted in massive proteinuria and renal failure in mice [39]. A previous study in mouse podocytes also indicated that the actin cytoskeleton could be a potential target for stabilizing cellular morphological changes, proteinuria, and renal function [39]. The interplay between pathways related to MLC kinase and phosphatase, Rho/ROCK, calcium, NO, VEGF, VEGFR, and ROS were prominently observed in fenestrated LSECs [14]; therefore, these are considered in the extended network.

We assume that the overall fenestration number depends on the actin cytoskeleton structure regulated by the proportion of stressed and relaxed actin fibers. The role of actin cytoskeleton as an essential structural and functional element that controls cell shape, cell motility, and adhesion has been demonstrated in different cell types [39, 40]. Myosins convert ATP to create a mechanical force on actin, which creates tension in the actomyosin cytoskeleton necessary for various functions [14]. Under a changing extracellular environment, actin structures are disassembled and remodeled to balance the structural and functional integrity. Thick actin stress fibers have been associated with elevated regions (raft regions) in endothelial cell monolayers. According to the sieve-raft hypothesis [12], these elevated regions have no fenestrations. It has been postulated that the regulation of fenestration size in LSECs is facilitated by the contraction or relaxation of the cytoskeleton surrounding the sieve plates [17, 25]. Fenestrations exist within the mesh-like structure comprised of actin [41]. The presence of stressed actin fibers leads to loss of fenestrations, whereas relaxed fibers promote fenestration formation. Thus, we consider both to determine the changes in the fenestration number. As the thick actin fibers around the mesh-like structure cause it to stretch, the fenestrations enlarge or increase in size [25]. On the other hand, when actin fibers relax and tension around the mesh-like structure loosens, fenestration size reduces. In the extended network, fenestration diameter is directly linked to pMLC protein, which influences both stressed and relaxed actin fibers (Fig 1).

### Logic-based network model development

The LBODEs modeling technique combines ordinary differential equations (ODEs) that are continuous functions of time with qualitative logic-based Boolean up- or down-regulation (i.e., activation or inhibition) using normalized Hill functions (saturating sigmoidal terms) for the logic-based modeling portion [26, 42]. The LBODEs framework allows predictions of network dynamics and is compatible with many analyses from nonlinear dynamics while requiring minimal knowledge of biochemical parameters [31, 43]. The LBODEs model uses a normalized-Hill function to define normalized species activity between 0 and 1 [26]. The LBODEs model parameters are categorized as reaction parameters—reaction weight (*W*), Hill coefficient (*n*), and half-effect (*EC*_50_)—and species parameters—maximum species activity (*y_max_*), initial species value (*y*_0_), and time constant (*τ*). Reaction weight regulates the strength of each network interaction, half-effect regulates the amount of input activity to achieve maximum output activity, and time constant regulates the time to activate or inhibit a network species. More detailed information on the LBODEs model structure, equations, and parameters can be found in our earlier work [26]. Our previous LBODEs model was thoroughly calibrated and validated against *in vitro* experimental datasets [26].

Here, we consider the expanded network (Fig 1) and *in vivo* data and scenarios. The optimal parameters in the previous LBODEs model are used in the extended model for the respective nodes. Due to limited information on model parameters for the additional nodes of the extended LBODEs model, we set these new parameters at their default values (*W_j_*= 1, *n_j_* = 1.4, 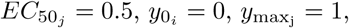 *τ_i_* = 1). The species and reaction parameters are listed in Tables A2 and A3 in S1 Appendix. The equations of the extended LBODEs model (Eqs A1–A35 in S1 Appendix) for the extended PSN (Fig 1) are generated automatically using Netflux, an open-source software package [42, 43]. The information about model parameters and reaction rules in the extended PSN (Tables A2 and A3 in S1 Appendix) is sufficient to generate the LBODEs using Netflux. Further, the LBODEs model is simulated in MATLAB after necessary modifications, as discussed below.

The cellular and protein expression responses to the glucose stimuli are governed by biochemical reactions or interactions in the network model. These reactions allow the transfer of information within or between cells, which can happen at a range of timescales (seconds to hours). The time constant parameter in the LBODEs model modulates the early or late activation of a protein in the network. The previous LBODEs model [26] was simulated and fitted for a short timescale (48 hours) suitable for *in vitro* studies. In the extended LBODEs model, time constant parameters for each protein come from a published timescale analysis [44]. The previous timescale analysis was performed for similar biochemical reactions, as seen in the extended network model, where time constant values were grouped by reaction type. The reaction types in the extended network are categorized into the following reaction types: ligand-receptor, transcription, translation, NF-κB activation, and signaling reactions. The time constants for the reaction types are as follows (Table A2 in S1 Appendix): ligand-receptor is 6 min, transcription is 88 hours, translation is 1.13 hours, NF-κB activation is 3.3 min, and signaling is 1 hour.

### Model calibration

For parameters not used at their default values, we calibrate the LBODEs model to *in vivo* data. These include the glucose dynamics and dynamic changes in fenestration structures.

#### Dynamic glucose stimulus

The extended model is stimulated by dynamic glucose levels in diabetic mice. A representative data set of *in vivo* glucose dynamics in diabetic mice [45] is used for the stimulus; glucose levels initially increase linearly from baseline and then fluctuate within a hyperglycemic range. Glucose concentration is fitted linearly (Eq 1) from 336–1008 hours (2–6 weeks) to data [45]:

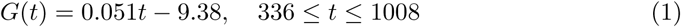

where *G*(*t*) is glucose concentration at time *t* in hours. After week 6, glucose is sampled from observed glucose concentrations in the diabetic mice population [10, 45] using step changes at each time point (i.e., concentration values are held constant during each interval between the measured data error bars). Glucose concentration is sampled in this stepwise fashion between upper and lower bounds at the following time points corresponding to published experimental measurement times [10, 45]: 7, 8, 9, 10, 11, 12, 14, 16, 18, and 20 weeks. The LBODEs model is simulated between 336–3360 hours (2–20 weeks).

Glucose input to the LBODEs model must be a normalized value on the order of 1, typically between 0 and 1. Therefore, a variable reaction weight 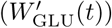 was calculated (Eq 2) from glucose concentration at each time by normalizing *G*(*t*) between 0 and 1:

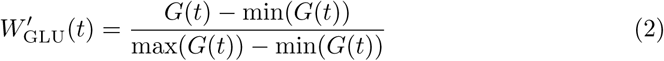

where min(*G*(*t*)) and max(*G*(*t*)) are the minimum and maximum concentrations of observed glucose, respectively. Due to a lack of reported data for glucose between 2 and 12 weeks in [10], min(*G*(*t*)) is set to baseline glucose concentration at 2 weeks as reported in Lee et al. [45]. Reported data by Finch et al. [10] is primarily used for model calibration. Therefore, max(*G*(*t*)) is set to the maximum value of the upper bound of observed glucose in Finch et al. [10]. Therefore, a normalized glucose activity of 1 corresponds to the maximum glucose concentration in mice reported in Finch et al. [10].

Normalized glucose activity (GLU) is used as the input to the extended LBODEs network and is calculated as

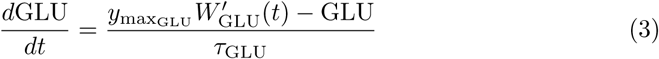

where 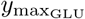 is the maximal glucose activity and is set to 1, 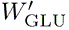 is the variable reaction weight for glucose from Eq 2, and *τ_GLU_*is the time constant for glucose and is set to 1.

#### Dynamic changes in fenestration structure

The additional species in the extended network are derived from published studies in other fenestrated endothelial cell types, as no experimental or animal models measure these proteins and validate these pathways in GECs under hyperglycemic conditions.

We simulate and validate the changes in fenestration number and diameter using a published *in vivo* study of GEC dysfunction in the advanced stage of DKD [10]. In our model, the fenestration number depends on the total activity of stressed and relaxed actin fiber. Relaxed actin fibers promote the formation of fenestrations at a rate *k*_form_, and stressed actin fibers reduce fenestrations in GECs at a rate *k*_loss_. Eq 4 is used to calculate the change in fenestration number (*y*_Number_) over time:

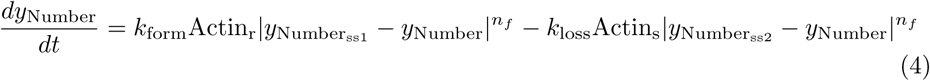

where *k*_form_ and *k*_loss_ are the rates of formation and loss of fenestrations, respectively. 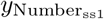 and 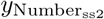 are the numbers of fenestrations at steady state for healthy control and diabetic mice, respectively. *n_f_*is the shape factor. Actin_s_ and Actin_r_ are stressed and relaxed states of actin fiber activity from Eqs A29 and A30 in S1 Appendix, respectively.

The predicted fenestration number is compared with measured fenestration density in GECs in mice, where density is the total number of fenestrations per unit length (pm) of the peripheral cytoplasm. The first term in Eq 4 defines a nonlinear increase in fenestration number compared to healthy mice. As a first step, only the first term in Eq 4 is considered, and three parameters are fit to fenestration density at 6, 10, 15, and 20 weeks in healthy mice [10]. Some necessary changes are made to the disease model to simulate fenestration formation in healthy cases. In healthy cases, we assume a balance between relaxed and stressed actin fiber activity. To observe balanced actin fiber activity, we switch to an activation interaction between ROCK and MLCP to promote the activation of relaxed actin fibers. We estimate *k*_form_, 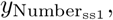 and *n_f_*using this methodology.

In the second step, we consider the disease model (Fig 1) and both terms in Eq 4. In diseased cases, the model promotes an imbalanced expression of relaxed and stressed fibers. The relaxed actin fibers are inactive in the diseased model mainly due to ROCK-mediated MLCP inhibition. The remaining parameters (*k*_loss_, 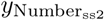) that define the decrease in fenestration number in the second term are fit using observed fenestration density at the same time points (6, 10, 15, and 20 weeks) in diabetic mice [10].

The phosphorylation of MLC protein (pMLC) (Eq A33 in S1 Appendix) is assumed to be the source of stress that increases fenestration diameter at a rate *k_s_* (Eq 5). This assumption is based on previous experimentation in fenestrated LSECs [14, 25]. Eq 5 is used to calculate the change in fenestration diameter (*y*_Diameter_) over time:

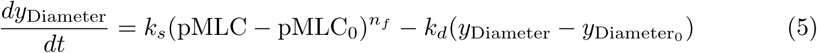

where *k_s_* and *k_d_* are the rates of increase in diameter (nm/hr) due to phosphorylation of MLC protein and restoration (1/hr) of diameter, respectively. pMLC_0_ and 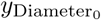 are the initial values of normalized pMLC protein and diameter at baseline in control subjects or healthy mice. The increased fenestration diameter is restored via an unknown restoring force at a rate of *k_d_*. Parameters *k_s_* and *k_d_* and time constant (*τ*_pMLC_) for pMLC activation are calibrated against observed data for fenestration width in GECs in diabetic mice in [10] using fmincon in MATLAB. The shape factor (*n_f_*) is set at the same value identified using Eq 4.

### Parameter estimation and uncertainty quantification

We perform a multi-start nonlinear least squares parameter estimation to estimate unknown parameter values in Eqs 4 and 5. We sample 25 parameter subsets from specified ranges for each parameter. *k*_form_, 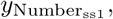 and *n_f_* are sampled in the ranges of [0.1,4] 1/hr, [6,8], and [2,5], respectively. *k*_loss_ and 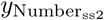 are sampled in the ranges of [1,5] 1/hr and [3,5], respectively. *k_s_*, *k_d_*, and *τ*_pMLC_ ae sampled in the ranges of [45, 75] nm/hr, [1,4] 1/hr, and [1,600] hr, respectively. The best-fit parameter values are the fitted parameter values that have the lowest sum of squared error (SSE) between prediction and data. These best-fit parameters are used for model prediction. The best-fit parameters are reported in Table 1 and Tables A2 and A3 in S1 Appendix.

**Table 1.**
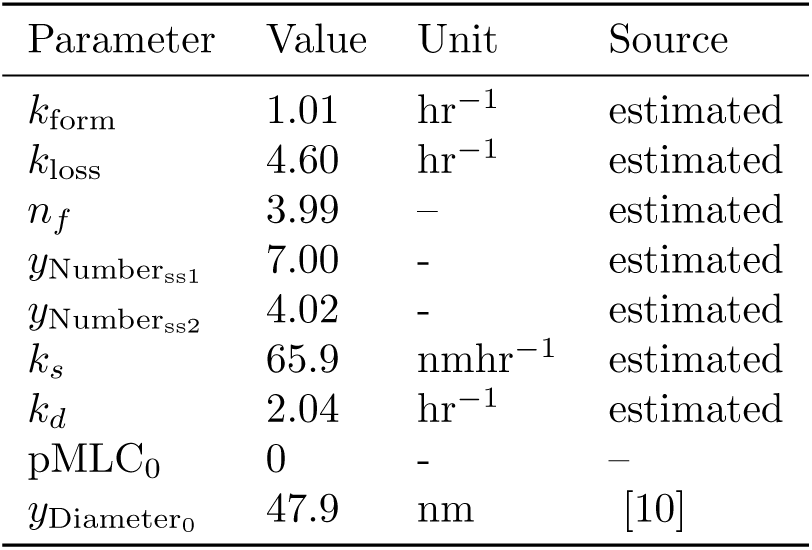
Parameters that modulate change in fenestration number and diameter.

We quantify the uncertainty in the model predictions for fenestration diameter and number using Monte Carlo ensemble simulation, a form of sampling-based uncertainty propagation [26, 46, 47]. To measure the parameters’ uncertainty, we use the function randsample in MATLAB to return a user-specified number of values sampled uniformly at random from the values in the vector population. We call randsample independently for each parameter to generate 1000 samples. The fitted parameter sets are labeled as an acceptable subset if the SSE for a given parameter subset is within 20% of the lowest SSE. For each parameter, we specify population as the values of that parameter in the acceptable parameters subset. The resulting 1000 values for each parameter are considered as the posterior distribution for the parameter. The posterior distributions of predictions for each output are used to calculate the 95% equal-tail credible interval at each time point, which defines the region where there is a 95% probability of containing a true estimate [26, 48, 49].

### Sensitivity analysis to determine targets for therapeutic interventions

*In silico* perturbation tests are simulated to identify the functional influence of each node under high glucose conditions. Similar *in silico* knockdown experiments and sensitivity analyses have been commonly used to study large network models [29, 31, 32]. Currently, no treatment strategies mitigate the loss of GEC fenestration in diabetic kidneys by leveraging precise mechanisms of action. *In silico* tests could identify disease intervention strategies and mechanisms to regulate endothelial dysfunction.

A recent study demonstrated several chemical agents’ dose-dependent role in regulating fenestration porosity and diameter of LSECs [25]. We test the qualitative effect of these chemical agents on fenestration structure in diseased GECs through our *in silico* framework. We perform *in silico* interventions to simulate the effects of the chemical agents on GEC fenestration structure using the extended LBODEs model. The effects of five chemical agents [25] are demonstrated through inhibitory actions on MLCK, calcium, ROCK, MLCP, and stressed actin fibers, respectively. *In silico* species inhibition is achieved by reducing 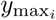 for respective species to 0. A statistical pairwise student’s t-test [50] is used to compare the differences in means between healthy group [10] and diseased groups with or without treatment using ttest2 in MATLAB. ttest2 uses observed or model-predicted values in each group to compute a mean for each group and compare them. Therefore, the extended LBODEs model is simulated 10 times with stepwise sampled glucose concentrations from a diabetic mice population. Each simulation generates different glucose concentrations between 6 and 20 weeks, representing uniform distributions of concentrations between the measured error ranges at each time interval to capture inter-subject variability. The model predicts fenestration number and diameter values at 20 weeks for each glucose simulation with or without an inhibitory treatment in diseased GECs. For each pairwise t-test [50], a significant difference between groups is reported when the calculated p-value is below the 0.001 significance level.

We also analyze the model under different simulated protein activity to screen for new and potential strategies for therapeutic intervention. We focus on two modes of therapeutic intervention: (1) protein knockdown or promotion and (2) reducing the strength of a reaction. We compare the sensitivity of each node in the network to perturbations in 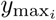 and *W_j_*. Knockdown or promotion of species is achieved by reducing or increasing 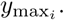 Reducing the reaction weight (*W_j_*) reduces the strength of the reaction. The local sensitivity index is defined as

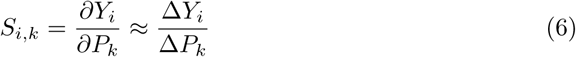

where *S_i,k_* is the sensitivity coefficient for a given output species *i* and parameter *k*, *Y_i_*is the output for species *i* at the optimal parameter values *P*, Δ*P_k_* is a small perturbation in parameter *k*, and Δ*Y_i_* = *Y_i_*(*P* + Δ*P_k_*) − *Y_i_*(*P*) is the change in output species *i* calculated at the perturbed parameter value. Eq 6 is used to calculate the local sensitivity index for fenestration number and fenestration diameter when each parameter is reduced by 100% from its optimal value (Tables A2 and A3 in S1 Appendix), simulating complete knockdown. We assess the sensitivity indices to help elucidate the functional effect of each protein or reaction on fenestrations. The magnitude of the sensitivity index suggests the degree of change in output (fenestration number or diameter). The positive sign of the sensitivity index suggests a decrease in output with a decrease in parameter value, and a negative sign suggests a decrease in output with an increase in the parameter value.

## Results

### Model simulations of disease onset and progression

The simulated glucose concentration and normalized glucose levels are shown in Fig 2. For weeks 2–6, Eq 1 was used for the glucose dynamics, and for weeks 6–20, the values were set at the observed mean of the measurements from Lee et al. [45] and Finch et al. [10]. Between measurement times, the simulated glucose concentration was held constant at the value from the previous time point. Then, the values were stepped piece-wise at each corresponding measurement time. The normalized results from transforming the values in Fig 2A using Eqs 2 and 3 are shown in Fig 2B.

**Fig 2.**
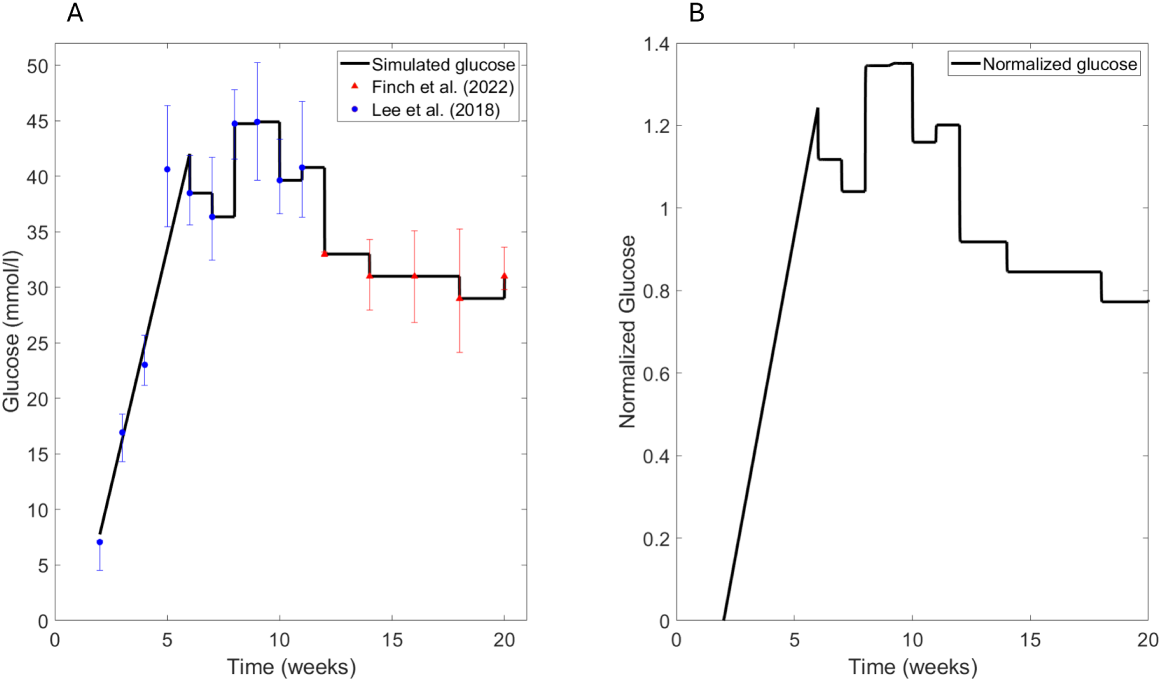
(A) Simulated glucose levels (black) over 20 weeks using the linear fit for weeks 2–6 from Eq 1 and piece-wise constant values of glucose at the observed data values in diabetic mice for weeks 6–20. Glucose concentration data values were obtained from Lee et al. [45] (blue) and Finch et al. [10] (red). (B) Normalized mean glucose levels using Eq 3.

Using the glucose dynamics shown in Fig 2A, we calibrated the predictions for the fenestration diameter (Eq 5) and fenestration number (Eq 5) to measured data from Finch et al. [10] (Fig 3). Fenestration diameter (Fig 3A) increased as early as 6 weeks, which directly correlated with full activation 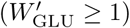 of normalized glucose around 6 weeks (Fig 3A) and subsequent signal transduction and protein activation in the network (Fig A1 in S1 Appendix). Similarly, we observed a loss in the fenestration number in agreement with the experimental data (Fig 3B).

**Fig 3.**
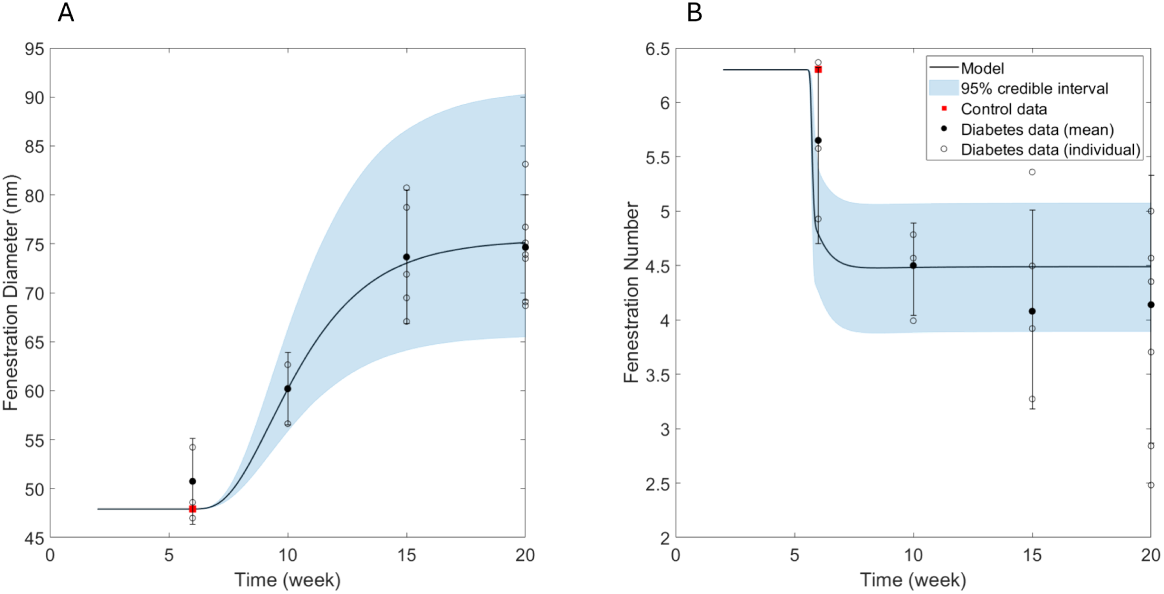
(A) Simulated fenestration diameter fitted against observed mean fenestration width (black circles) in diabetic mice [10]. (B) Simulated fenestration number fitted against observed mean fenestration density (black circles) in diabetic mice [10]. Blue shaded regions show the 95% credible intervals of the predictions. Individual fenestration width and density are also reported for each diabetic mouse (open circles). Initial values for fenestration diameter and number are assumed to be the same as baseline control data values of fenestration width and density (red squares) for healthy mice in Finch et al. [10].

### Glucose-mediated effect on fenestration loss and diameter

The extended model was also helpful in making model-informed predictions about potential disease interventions and treatments. In the previous section, we simulated the species activity (Fig A1 in S1 Appendix) at the simulated glucose levels sampled at the observed mean of the data in diabetic mice [10]. Here, we explored the effects of between-subject variability in glucose concentration on the changes in species activity and fenestration structure by running 25 sampled glucose profiles (Fig 4A). A uniform distribution of glucose profiles between each set of experimentally determined error bars spread the samples over a broader range than the experimental normal distribution, where values are closer to the mean. Uniform sampling of glucose concentration allowed us to explore greater inter-subject variability. We observed no significant variations in fenestration dynamics and most species due to variations in glucose concentrations across subjects (Fig A2 in S1 Appendix). AGE, RAGE, and NADPH node activity varied with glucose variations significantly (Fig A2 in S1 Appendix). The variability in glucose concentration was mainly observed in the hyperglycemic range and indicated impaired glucose tolerance.

**Fig 4.**
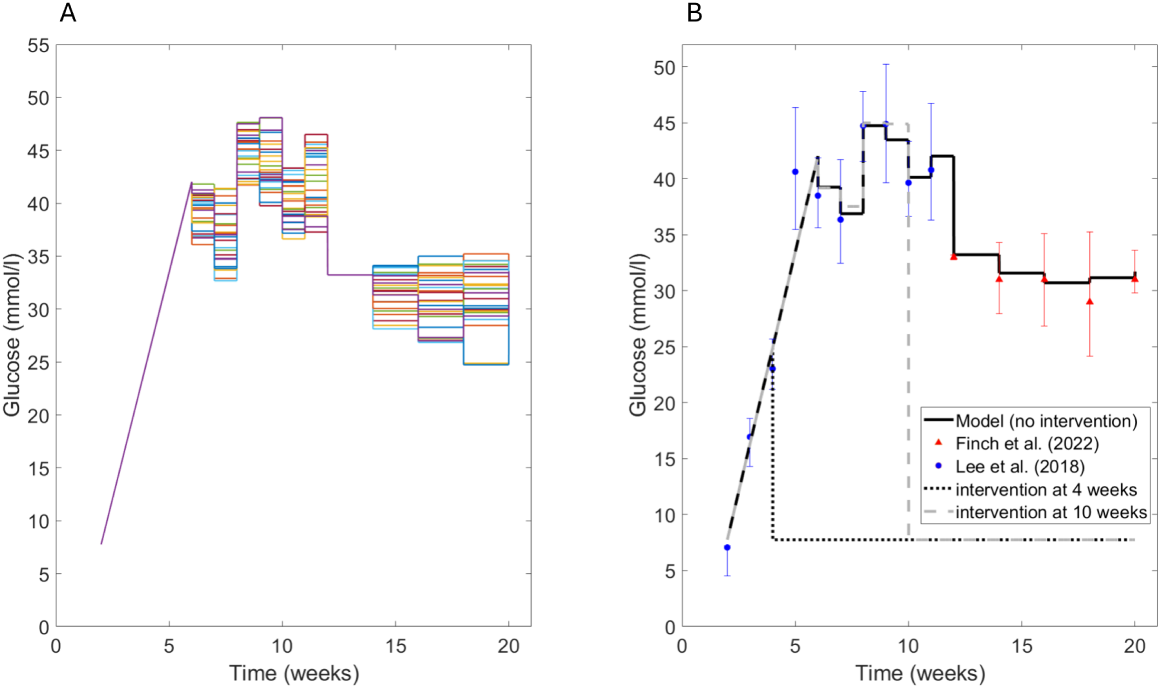
(A) Simulated glucose concentration profiles sampled uniformly 25 times. (B) Simulated average glucose concentration (solid black lines) without intervention, *in silico* glucose control at 4 weeks (dotted black line) and 10 weeks (dashed gray line) are shown. Glucose concentration data values were obtained from Lee et al. [45] (blue) and Finch et al. [10] (red).

In mice carrying the diabetes mutation (leptin-deficient), a manifestation of the diabetic syndrome depends on genetic background. In these mice used as the reference for this study, male mice developed diabetes around six weeks of age [51–53]. In Fig 4, glucose levels were controlled *in silico* at time points before and after the mice reached the diabetic state. Fig 5 shows the changes in species activity relative to baseline upon glucose intervention (Fig 4) at different time points between 2–20 weeks.

**Fig 5.**
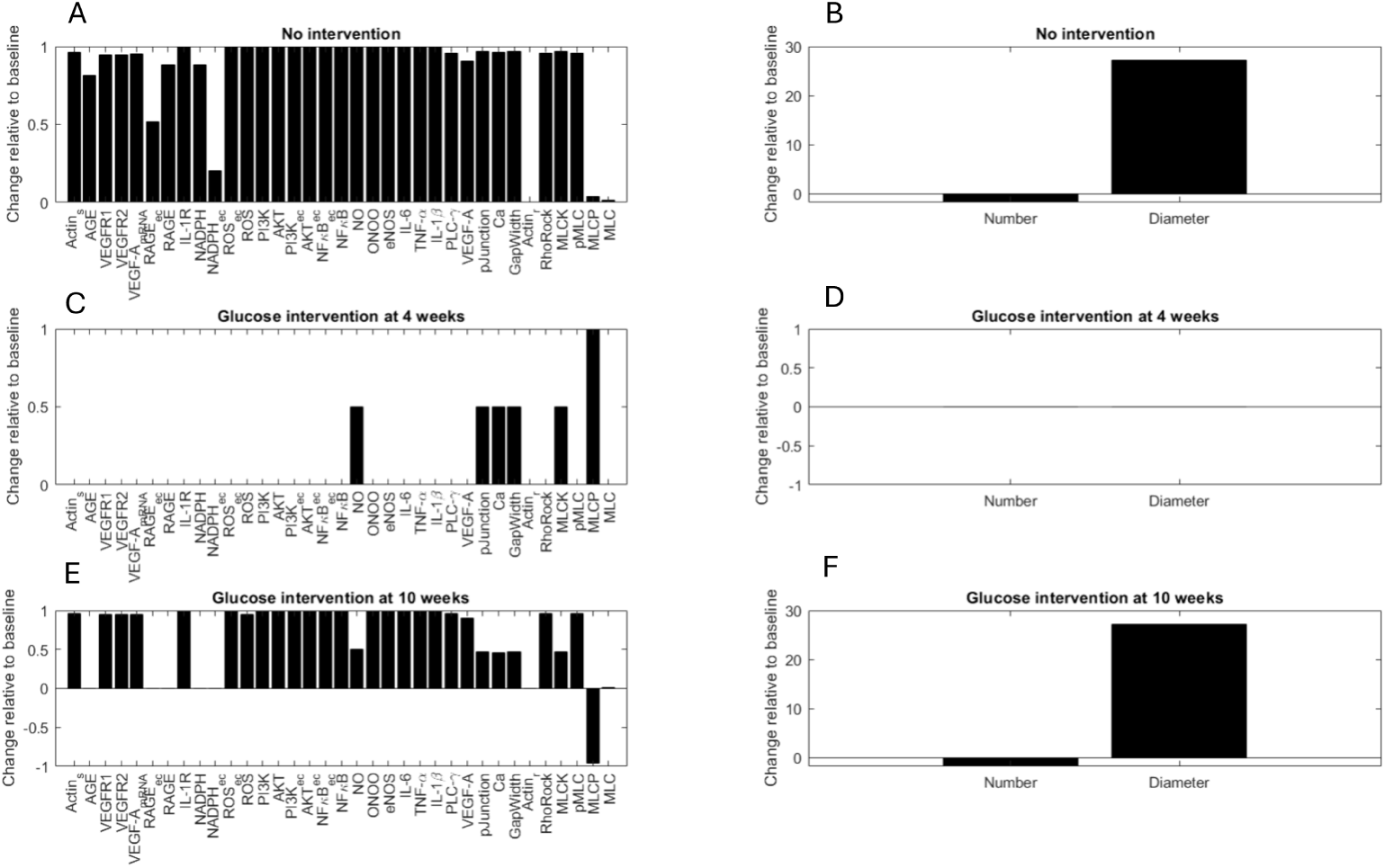
(Predicted change in (A) activity and (B) fenestration number and diameter from baseline without glucose intervention. Predicted change in (C) activity and (D) fenestration number and diameter from baseline upon glucose intervention at 4 weeks. Predicted change in (E) activity and (F) fenestration number and diameter from baseline upon glucose intervention at 10 weeks.

The model predicted that glucose intervention at 4 weeks was most influential in reversing upregulated protein expression and controlling changes in fenestration structure (Fig 5). Glucose control at 4 weeks maintained balance in NO and calcium activity, which may be relevant in regulating downstream signaling dysfunction. MLCK protein levels decreased, and MLCP protein levels increased considerably upon glucose intervention. On the contrary, a glucose intervention at 10 weeks was ineffective in controlling fenestration dynamics; however, it controlled the upregulation of AGE, RAGE, and NADPH activity in macrophages and GECs. The normalized glucose levels reached their maximal activity by 5 weeks, which led to a self-sustained maximal activation of species in the network. The minimum activity of most species was equal to or above their *EC*_50_ by 5 weeks. The positive feedback loops in the network also regulated the self-sustained maximum activity of downstream species. This resulted in consistently dysregulated signaling, loss of fenestrations, and increased fenestration diameter. Together, these results suggest that glucose control may be an effective strategy for controlling and mediating GEC activation in the early stages of DKD. Yet, it cannot be used as the only strategy to modulate complex signals and pathways that regulate fenestrations in the later stages of DKD.

### Proposed treatment and intervention strategies

We compared the effects of KN93, ML-7, Y27632, Calyculin A, and Cytochalasin B on their respective targets that we assumed were shared between LSECs and GECs. KN93, ML-7, Y27632, Calyculin A, and Cytochalasin B inhibit calcium, MLCK, ROCK, MLCP, and stressed actin fibers, respectively, in fenestrated endothelial cells. Fig 6 compares the simulated fenestration number and diameter in diseased GECs at 20 weeks without treatment, with *in silico* treatment with these chemical agents, and reference values observed in healthy mice. The error bars for predicted fenestration number and diameter were small and not included with the results (Fig 6).

**Fig 6.**
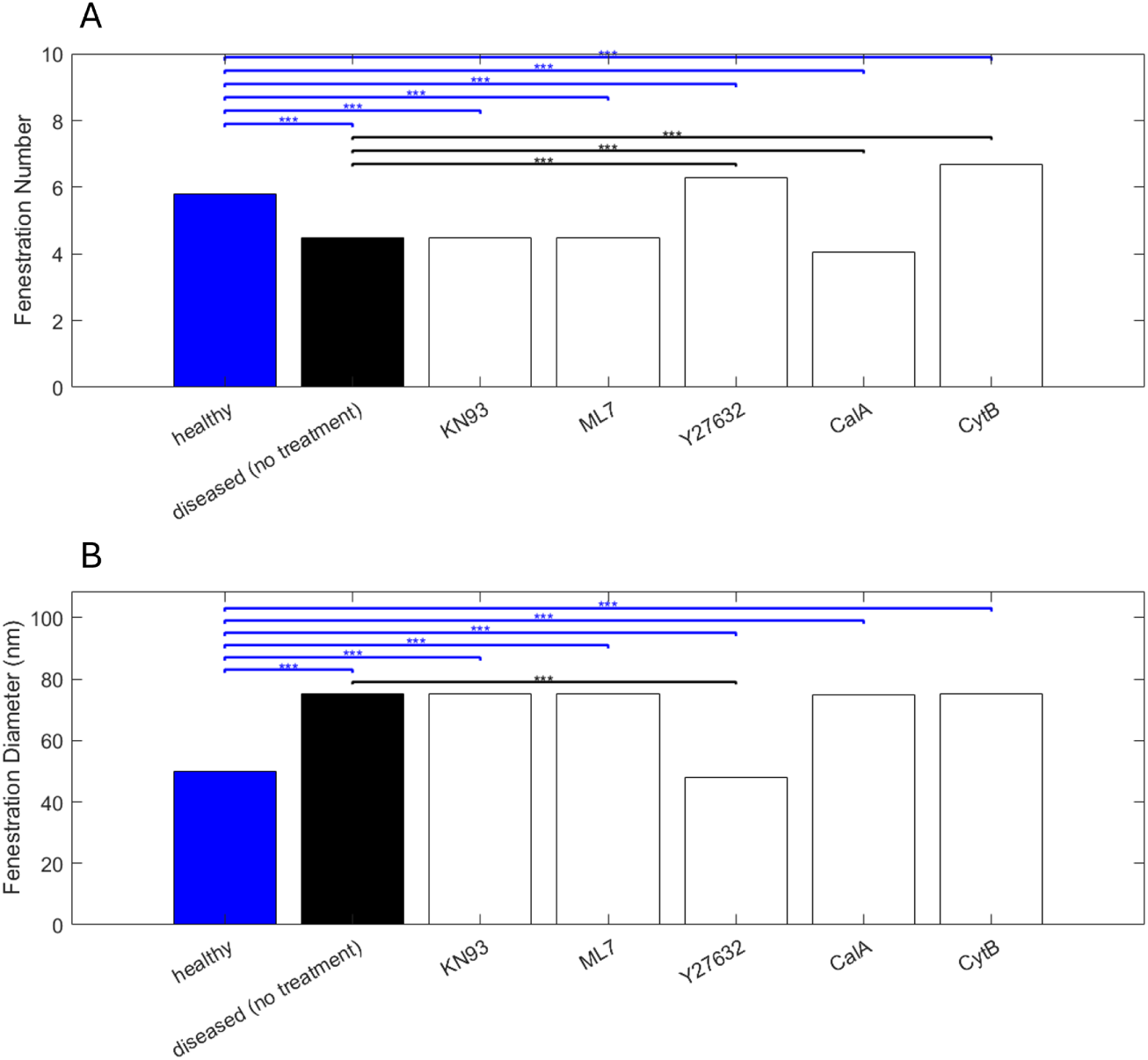
Bar plots comparing the simulated (A) fenestration number and (B) diameter at 20 weeks in diseased mice before (black) and after treatment with respective chemical agents (white). Observed number and diameter in healthy mice from Finch et al. [10] are also shown (blue bars). ***: p-value*<* 0.001 for t-test comparison between healthy mice and model predictions in diseased GECs with or without treatment. ***: value*<* 0.001 for t-test comparison between model predictions in diseased GECs without treatment and model predictions with treatment.

Fenestration number significantly increased upon treatment with Y27632 and Cytochalasin B, which inhibited ROCK protein and stressed actin fibers, respectively, compared to both diseased and healthy cases (Fig 6). The predicted fenestration number decreased further compared to both diseased and healthy cases when MLCP protein was inhibited by Calyculin A. No significant changes were observed in fenestration number upon treatment with KN93 and ML-7 compared to untreated diseased GECs. However, treatment with KN93 and ML-7 had a substantial effect in decreasing the fenestration number when compared to healthy GECs. Fenestration diameter decreased significantly upon inhibition of ROCK protein using Y27632. KN93, ML-7, Calyculin A, and Cytochalasin B were ineffective in regulating the diameter of diseased GECs. However, compared to healthy GECS, each *in silico* inhibition significantly affected diameter in diseased GECs.

Based on the sensitivity analysis, we found that the following species were most effective in controlling fenestration diameter when controlled early in diabetic mice (before 5 weeks) (Fig A4 in S1 Appendix): glucose, AGE, VEGFR-2, VEGF-A (mRNA), RAGE*_ec_*, IL-1R, NADPH*_ec_*, ROS*_ec_*, RAGE*_ec_*, NFκB*_ec_*, NFκB, IL-1β, VEGF-A, ROCK, and pMLC. Among the influential reactions that positively correlated with fenestration diameter, the following reactions decreased fenestration diameter (Fig A4 in S1 Appendix): IL-1β ⇒ IL-1R, GLU ⇒ AGE, VEGF-A ⇒ VEGFR2, AGE ⇒ RAGE*_ec_*, ROS*_ec_* ⇒ NFκB*_ec_*, RhoROCK ⇒ pMLC, VEGFR2 ⇒ RhoROCK, NFκB*_ec_* ⇒ IL-1β, NFκB ⇒ VEGF-A (mRNA), VEGF-A (mRNA) ⇒ VEGF-A, NADPH*_ec_* ⇒ ROS, RAGE*_ec_* ⇒ NADPH*_ec_*. MLCP, MLC, and relaxed actin fibers negatively correlated with the fenestration number, as evident from the sensitivity analysis (Fig A3 in S1 Appendix). Glucose, AGE, VEGFR-2, VEGF-A (mRNA), RAGE*_ec_*, IL-1R, NADPH*_ec_*, ROS*_ec_*, RAGE*_ec_*, NFκB*_ec_*, NFκB, IL-1β, VEGF-A, ROCK, pMLC, and stressed actin fibers had a positive correlation with fenestration number (Fig A3 in S1 Appendix). We observed only minor differences in sensitivity indices for fenestration number upon local perturbations in reaction weight (Fig A3 in S1 Appendix).

In most studies, it was reported that diabetes may develop and progress into diabetic kidney disease around 10–12 weeks in diabetic mice [45]. Therefore, it may be relevant to understand when to intervene as DKD develops and progresses in these subjects and to effectively regulate fenestration number and diameter before damage worsens. To study this *in silico*, we inhibited influential species and interactions identified by the sensitivity analysis by 50% at 8, 10, and 20 weeks to compare the effects on fenestration structure during pre-DKD, early-DKD, and late-DKD stages in diabetic mice, respectively. The dynamic effects of inhibiting these species and interactions on fenestration structure during the early and late stages of DKD in mice were simulated. For an *in silico* knockdown of influential species at 8 weeks in diabetic mice, an increase in fenestration number was mostly achieved with just 50% inhibition of stressed actin fibers, VEGF-A (mRNA), VEGF-A, ROCK, and NFκB (Fig 7A). A slight increase in fenestration number was achieved by promoting the activity of MLC, MLCP, and relaxed actin fibers (results not shown). Upon reducing the strength of interaction between VEGFR2 and RhoROCK protein, and between VEGF-A (mRNA) and VEGFR2, a significant increase in fenestration number was observed (Fig 7B). 50% inhibition of ROCK, VEGF-A, VEGFR2, pMLC, and NFκB restored fenestration diameter significantly (Fig 7C). Reducing the strengths of interaction between ROCK and pMLC and between NFκB and VEGF-A (mRNA) were most effective in restoring fenestration diameter (Fig 7D). The inhibition time did not significantly affect the time to recover fenestration number or diameter. We observed a similar effect of inhibited species or reduced reaction strength on fenestration number and diameter at 10 weeks (Fig A5 in S1 Appendix) and 20 weeks (Fig A6 in S1 Appendix). Glucose, AGE, RAGEec, NADPHec, ROSec, NFκBec inhibition did not affect fenestrations at 8 weeks (data not shown), 10 weeks (Fig A5 in S1 Appendix), or 20 weeks (Fig A6 in S1 Appendix).

**Fig 7.**
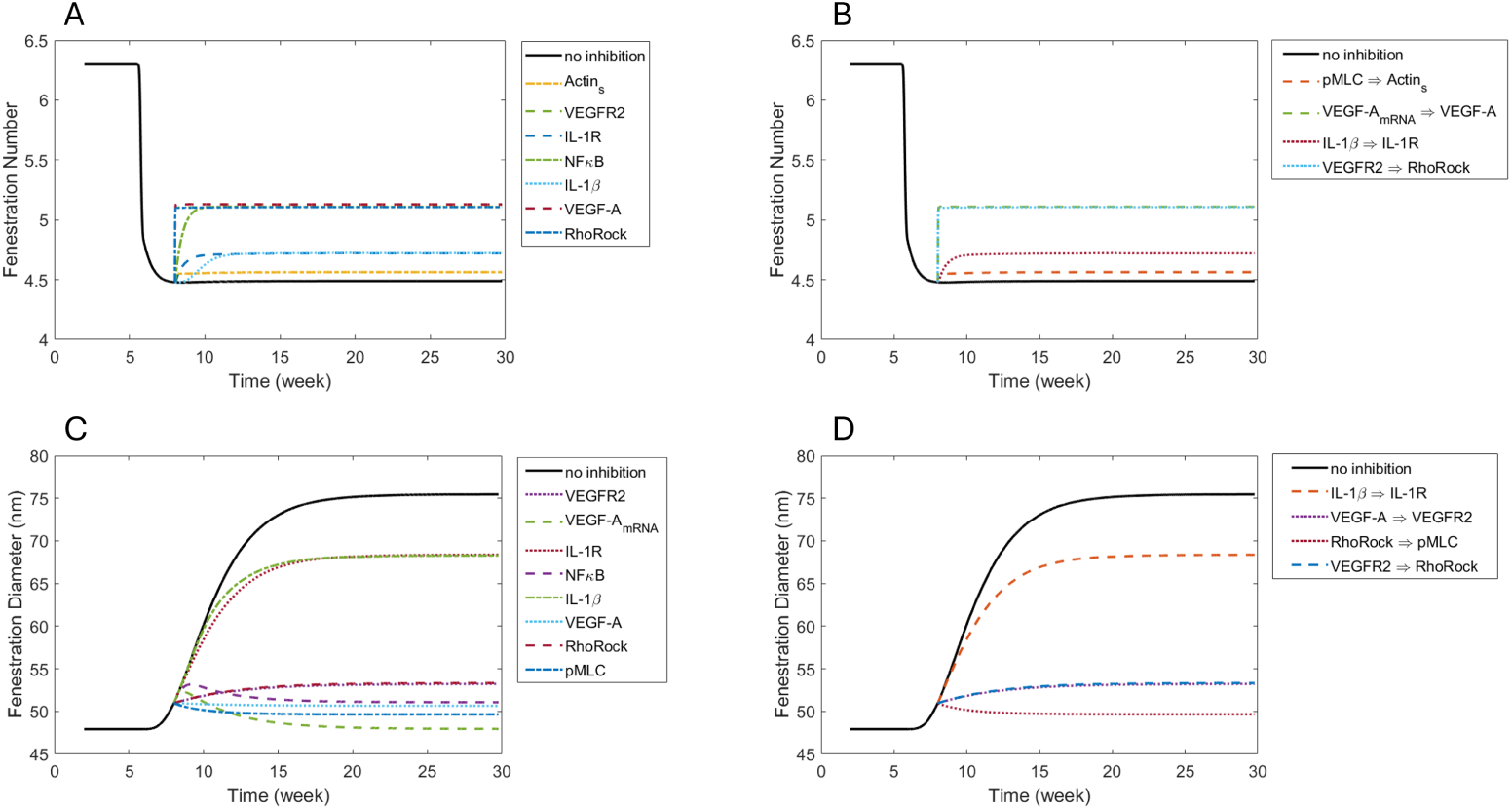
Effects at 8 weeks of 50% inhibition of sensitive (A) species and (B) reactions on fenestration number and of sensitive species (C) and (D) reactions on fenestration diameter.

## Discussion

Glomerular endothelial activation and dysfunction are early signs of DKD development and progression. Alterations in the size and density of GEC fenestrations have been recently associated with the disruption in glomerular filtration and the progression of diabetic kidney disease [10]. Currently, significant barriers impede understanding the cellular mechanisms that regulate GEC fenestrations, which include lack of ideal *in vitro* models, fenestration loss in culture, and inconsistencies between *in vitro* and *in vivo* findings [9]. The small size of endothelial cell fenestrations is beyond the limit of resolution of light microscopy and needs newer and advanced technology to be studied accurately. Hence, there is a lack of high-quality quantitative data on transient changes in fenestration size and density in diseased states. It is beneficial to leverage mathematical modeling to study GEC fenestrations and explore signaling drivers that affect their size and density. In this study, we presented an extension to our previously developed protein-protein signaling network model and studied the effects of glucose-mediated signaling dysregulation on structural changes in fenestrated GECs using an *in silico* approach.

Signaling drivers and mechanisms were derived from fenestrated LSECs and considered in the presented model. The interplays between pathways modulating MLCK, Rho/ROCK, calcium, NO/eNOS, VEGF/VEGFR, ROS, and actin structure were crucial in understanding fenestration dynamics. Under a changing extracellular environment, actin structures are disassembled and remodeled to balance the structural and functional integrity of the fenestrations [14]. Stresses within the actin structure are correlated with GECs’ fenestration density, and MLC protein phosphorylation is associated with the enlargement of fenestrations [25]. We demonstrated that glucose intolerance and hyperglycemia in diabetic mice resulted in the loss of about half of the fenestrations and a 70% increase in fenestration size in GECs from baseline. Previous studies also reported changes in fenestration in the glomeruli in patients diagnosed with diabetic nephropathy (DN) [10]. Fenestration loss and alterations in fenestration width for DN patients [10] were also quantitatively similar to the predicted changes in diabetic mice.

We showed that gradual increases in glucose levels with the development of diabetic conditions were correlated with dysregulated species and changes in the size and density of fenestrations. We observed that between-subject variability in glucose concentration in diabetic mice had minimal or no effect on normalized protein activity in macrophages or GECs. Glucose-mediated signaling dysregulation and autocrine inflammatory feedback were mathematically related to loss of fenestration number and increased diameter.

Previously published logic-gated network models [31, 54] have drawn mathematical relations between protein signaling and normalized changes in the cell area using LBODEs. Compared to previous models, our model related normalized signaling activity to actual changes in fenestration structure in diabetic mice.

We observed that glycemic control in the early stages was most effective in reducing upregulated species and modulating fenestration loss. After glucose concentration reached 25 mmol/l, often considered high glucose in mice, glucose control was ineffective for controlling GEC activation, signaling dysregulation, and structure in the later stages of DKD development. We propose that other disease intervention strategies besides glucose control are essential in regulating the complex downstream signals and pathways that regulate fenestrations in diseased GECs. This was seen through the predicted imbalanced activity of proteins in the MLC phosphorylation cycle (pMLC, MLCP, MLCK, and MLC) after glucose control (Fig 5).

Using sensitivity analyses, selective *in silico* knockdown of species provided alternative strategies for disease intervention. Strategies to inhibit stressed actin fibers, VEGF-A (mRNA), VEGF-A, ROCK, and NFκB were predicted to effectively recover the loss in fenestrations in diseased GECs. ROCK, VEGF-A, VEGFR2, pMLC, and NFκB protein inhibition were predicted to be promising strategies for controlling fenestration diameter in diseased GECs.

Chemical agents that inhibit ROCK protein and actin depolymerization agents like cytochalasin B have also been shown to increase porosity in cultured LSECs in a few minutes to hours [16]. However, previous experiments in LSECs [25] compared the effect of targeted inhibition using different chemical agents to healthy LSECs. A recent experimental study showed that treatment with cytochalasin B increased fenestration number and diameter in human renal GECs [2]. Using the extended model, we observed a qualitatively similar regulatory effect of Y27632 on ROCK inhibition and cytochalasin B on stressed actin inhibition in GECs (Fig 6). No published studies have tested the effects of targeted inhibition of MLCK, ROCK, and MLCP proteins in GECs affected by diabetes. The combined effects of ROCK inhibitors and depolymerization agents may be worth exploring to restore both porosity and fenestration diameter in diseased GECs. The model-based predictions also suggested that a balance in MLCP and MLCK proteins may be crucial in regulating fenestration diameter and number (fig 5). A relatively higher MLCK protein than MLCP protein was observed during fenestration structure damage despite glucose intervention at 10 weeks Fig 5. A balanced MLCP and MLCK expression should enhance fenestration formation (Fig A3 in S1 Appendix). At the same time, MLCP and MLCK perturbations had potentially no or minimal change in fenestration diameter as seen from the sensitivity analysis (Fig A4). The interdependency on the MLCP and MLCK protein balance is essential due to the positive feedback loops in the MLC phosphorylation cycle. Both MLCK and MLCP proteins regulated the fenestration diameter and porosity in LSECs [25]. Counterintuitive to the model predictions, inhibition of MLCK and MLCP by chemical agents like ML7 and Calcyculin A, respectively, resulted in fenestration loss and an increase in fenestration diameter (Table A4 in S1 Appendix). The model suggests potential strategies for disease intervention that can complement established methods once validated. Often, clinical biomarkers of kidney disease prediction and progression risk, such as serum creatinine and albuminuria, only show alterations relatively late in the disease process and, thus, may not be suitable for early disease diagnosis [55]. Therefore, the proposed mechanisms support our understanding of new disease biomarkers that are more closely related to histological changes in early DKD development and progression, and when validated, they can potentially improve the early diagnosis and clinical management of the disease.

We acknowledge the limitations of the model and the potential for further improvement. The protein activity is quantified as a fractional activation or inhibition rather than absolute quantities, such as concentration, and is limited to analyses with normalized chemical species levels. However, the predicted normalized activity of species can be transformed to their actual values if needed. Here, the fenestration number was considered as a continuum, representing an average value across all regions, not a discrete integer number of fenestrations. The presented LBODEs framework represents all protein-protein interactions using an activating or inhibiting Hill function, but other mathematical functions could be used. The LBODEs framework was useful in identifying potential disease intervention strategies or relevant mechanisms of action that can be used to design fit-for-purpose and less complex quantitative models with limited parametrization. The calibration and validation of the logic-based modeling framework was limited to *in vitro* and *in vivo* mouse studies. The model accuracy can be further improved through validation with other *in vitro* or *in vivo* data as available or through emerging data from single-cell analyses. There is a lack of knowledge related to the fundamental biology of the regulating GECs. More advanced imaging techniques known as super-resolution microscopy may offer the potential to facilitate the accurate measurement of fenestrations in GECs [9]. The proposed mechanistic interactions used to study fenestration dynamics in diabetic mice are also relevant in predicting changes in clinical population as a similar quantitative effect on fenestrations in DN patients was observed [10].

## Conclusion

In this study, we used a previously developed LBODEs model of protein-protein interaction and extended it to study the development and progression of DKD. The LBODEs network model predicted the effects of high glucose and inflammation on macrophage phenotypic changes, GEC activation, and signaling dysregulation in diabetic mice. Further, the extended LBODEs model related the signaling dysregulation with histological changes in GEC fenestrations, and mechanistic relationships were validated using *in vivo* mice data. Through *in silico* targeted inhibition, we identified the effective response time and validated mechanisms of action and effect on fenestrations through glucose control, species or pathway inhibition, or known chemical agents tested in other fenestrated cell types. We identified that disease intervention strategies besides glucose control are essential in regulating the complex downstream signals and pathways regulating fenestrations in diseased GECs. Network species, such as ROCK, VEGF-A, VEGFR2, pMLC, and NFκB, restored fenestration structure and density significantly. Reducing the interaction strength between ROCK and pMLC and between NFκB and VEGF-A (mRNA) were most effective in restoring fenestration diameter. The novel logic-based network model helped to quantify the crosstalk between macrophages and GECs in the early through late stages of DKD. The proposed mechanisms support our understanding of new disease biomarkers or pathways more closely related to histological changes in DKD development and progression. The proposed model could be integrated with more complex models to better predict disease progression and identify early biomarkers for DKD, enhancing clinical management and intervention strategies.

## Data Availability Statement

There are no primary data in the paper. The supporting MATLAB code, including parameter values, scripts for plotting, data, and documentation, can be accessed through an open-source GitHub repository available at https://github.com/ashleefv/KidneyImmuneLBM/tree/master/LBODE_extended_model.

## Supporting Information

**S1 Appendix.** Supplementary equations (Eqs A1–A35), supplementary tables (Tables A1–A4), and supplementary figures (Figs A1–A6).

## Acknowledgments

We thank Ford Versypt Lab members and Krutika Patidar’s Ph.D. committee members, Dr. Rudiyanto Gunawan and Dr. David Kofke, for their thorough feedback.

## Author Contributions

**Conceptualization:** Krutika Patidar, Ashlee N. Ford Versypt.

**Formal analysis:** Krutika Patidar, Ashlee N. Ford Versypt.

**Funding acquisition:** Ashlee N. Ford Versypt.

**Investigation:** Krutika Patidar, Ashlee N. Ford Versypt.

**Methodology:** Krutika Patidar, Ashlee N. Ford Versypt.

**Project administration:** Ashlee N. Ford Versypt.

**Software:** Krutika Patidar, Ashlee N. Ford Versypt.

**Visualization:** Krutika Patidar.

**Writing - original draft:** Krutika Patidar.

**Writing - review & editing:** Krutika Patidar, Ashlee N. Ford Versypt.

## References

1. Jourde-Chiche N, Fakhouri F, Dou L, Bellien J, Burtey S, Frimat M, et al. Endothelium Structure and Function in Kidney Health and Disease. Nat Rev Nephrol. 2019;15:87–108. doi:10.1038/s41581-018-0098-z.

2. Meijer EM, van Dijk CGM, Giles R, Gijsen K, Chrifi I, Verhaar MC, et al. Induction of Fenestrae in Human Induced Pluripotent Stem Cell-Derived Endothelial Cells for Disease Modeling. Tissue Eng Part A. 2024;30:168–180. doi:10.1089/ten.TEA.2023.0236.

3. Jiang S, Luo M, Bai X, Nie P, Zhu Y, Cai H, et al. Cellular crosstalk of glomerular endothelial cells and podocytes in diabetic kidney disease. J Cell Commun Signal. 2022;16:313–331. doi:10.1007/s12079-021-00664-w.

4. Albrecht M, Sticht C, Wagner T, Hettler S, Delatorre C, Qiu J, et al. The crosstalk between glomerular endothelial cells and podocytes controls their responses to metabolic stimuli in diabetic nephropathy. Sci Rep. 2023;13. doi:10.1038/s41598-023-45139-7.

5. Satchell SC, Braet F. Glomerular endothelial cell fenestrations: an integral component of the glomerular filtration barrier. Am J Physiol Renal Physiol. 2009;296:F947–F956. doi:10.1152/ajprenal.90601.2008.

6. Li T, Shen K, Li J, Leung SWS, Zhu T, Shi Y. Glomerular Endothelial Cells Are the Coordinator in the Development of Diabetic Nephropathy. Front Med. 2021;8:655639. doi:10.3389/fmed.2021.655639.

7. Deen WM. What determines glomerular capillary permeability? J Clin Invest. 2004;114:1412–1414. doi:10.1172/JCI23577.

8. Deen W, Lazzara M, Myers B. Structural determinants of glomerular permeability. Am J Physiol Renal Physiol. 2001;281:F579–F596. doi:10.1152/ajprenal.2001.281.4.F579.

9. Finch NC, Neal CR, Welsh GI, Foster RR, Satchell SC. The unique structural and functional characteristics of glomerular endothelial cell fenestrations and their potential as a therapeutic target in kidney disease. Am J Physiol Renal Physiol. 2023;325:F465–F478. doi:10.1152/ajprenal.00036.2023.

10. Finch NC, Fawaz SS, Neal CR, Butler MJ, Lee VK, Salmon AJ, et al. Reduced glomerular filtration in diabetes is attributable to loss of density and increased resistance of glomerular endothelial cell fenestrations. J Am Soc Nephrol. 2022;33:1120–1136. doi:10.1681/ASN.2021030294.

11. Denzer L, Muranyi W, Schroten H, Schwerk C. The Role of PLVAP in Endothelial Cells. Cell Tissue Res. 2023;392:393–412. doi:10.1007/s00441-023-03741-1.

12. Cogger VC, Roessner U, Warren A, Fraser R, Le Couteur DG. A Sieve-Raft Hypothesis for the Regulation of Endothelial Fenestrations. Comput Struct Biotechnol J. 2013;8:e201308003. doi:10.5936/csbj.201308003.

13. Zapotoczny B, Braet F, Kus E, Ginda-Mäkelä K, Klejevskaja B, Campagna R, et al. Actin-spectrin scaffold supports open fenestrae in liver sinusoidal endothelial cells. Traffic. 2019;20:932–942. doi:10.1111/tra.12700.

14. Szafranska K, Kruse LD, Holte CF, McCourt P, Zapotoczny B. The whole story about fenestrations in LSEC. Front Physiol. 2021;12:735573. doi:10.3389/fphys.2021.735573.

15. Cooper GM. Structure and Organization of Actin Filaments. In: The Cell: A Molecular Approach. Sinauer Associates; 2000.

16. Zapotoczny B, Szafranska K, Kus E, Braet F, Wisse E, Chlopicki S, et al. Tracking Fenestrae Dynamics in Live Murine Liver Sinusoidal Endothelial Cells. Hepatology. 2019;69:876–888. doi:10.1002/hep.30232.

17. Braet F, Zanger RD, CrabbÉ E, Wisse E. New Observations on Cytoskeleton and Fenestrae in Isolated Rat Liver Sinusoidal Endothelial Cells. J Gastroenterol Hepatol. 1995;10:S3–S7. doi:10.1111/j.1440-1746.1995.tb01792.x.

18. Paniagua OA, Bryant MB, Panza JA. Role of Endothelial Nitric Oxide in Shear Stress–Induced Vasodilation of Human Microvasculature. Circulation. 2001;103:1752–1758. doi:10.1161/01.CIR.103.13.1752.

19. Locatelli M, Zoja C, Conti S, Cerullo D, Corna D, Rottoli D, et al. Empagliflozin protects glomerular endothelial cell architecture in experimental diabetes through the VEGF-A/caveolin-1/PV-1 signaling pathway. J Pathol. 2022;256:468–479. doi:10.1002/path.5862.

20. Selby NM, Taal MW. An updated overview of diabetic nephropathy: Diagnosis, prognosis, treatment goals and latest guidelines. Diabetes Obes Metab. 2020;22:3–15. doi:10.1111/dom.14007.

21. Fu J, Lee K, Chuang PY, Liu Z, He JC. Glomerular Endothelial Cell Injury and Cross talk in Diabetic Kidney Disease. Am J Physiol Renal Physiol. 2015;308:F287–F297. doi:10.1152/ajprenal.00533.2014.

22. Bates DO. Vascular endothelial growth factors and vascular permeability. Cardiovasc Res. 2010;87:262–271. doi:10.1093/cvr/cvq105.

23. Tiruppathi C, Minshall RD, Paria BC, Vogel SM, Malik AB. Role of Ca2+ Signaling in the Regulation of Endothelial Permeability. Vasc Pharmacol. 2002;39:173–185. doi:10.1016/S1537-1891(03)00007-7.

24. Yokomori H, Yoshimura K, Funakoshi S, Nagai T, Fujimaki K, Nomura M, et al. Rho Modulates Hepatic Sinusoidal Endothelial Fenestrae Via Regulation of the Actin Cytoskeleton in Rat Endothelial Cells. Lab Invest. 2004;84:857–864. doi:10.1038/labinvest.3700114.

25. Zapotoczny B, Szafranska K, Lekka M, Ahluwalia BS, McCourt P. Tuning of liver sieve: the interplay between actin and myosin regulatory light chain regulates fenestration size and number in murine liver sinusoidal endothelial cells. Int J Mol Sci. 2022;23:9850. doi:10.3390/ijms23179850.

26. Patidar K, Ford Versypt AN. Logic-Based Modeling of Inflammatory Macrophage Crosstalk with Glomerular Endothelial Cells in Diabetic Kidney Disease. bioRxiv. 2023;Preprint:2023.04.04.535594. doi:10.1101/2023.04.04.535594.

27. Wu Q, Finley S. Mathematical model predicts effective strategies to inhibit VEGF-eNOS signaling. J Clin Med. 2020;9:1255. doi:10.3390/jcm9051255.

28. Weinstein N, Mendoza L, Gitler I, Klapp J. A network model to explore the effect of the micro-environment on endothelial cell behavior during angiogenesis. Front Physiol. 2017;8:960. doi:10.3389/fphys.2017.00960.

29. Liu X, Zhang J, Zeigler AC, Nelson AR, Lindsey ML, Saucerman JJ. Network Analysis Reveals a Distinct Axis of Macrophage Activation in Response to Conflicting Inflammatory Cues. J Immunol. 2021;206:883–891. doi:10.4049/jimmunol.1901444.

30. Zhao C, Medeiros TX, Sové RJ, Annex BH, Popel AS. A Data-Driven Computational Model Enables Integrative and Mechanistic Characterization of Dynamic Macrophage Polarization. iScience. 2021;24:102112. doi:10.1016/j.isci.2021.102112.

31. Ryall KA, Holland DO, Delaney KA, Kraeutler MJ, Parker AJ, Saucerman JJ. Network Reconstruction and Systems Analysis of Cardiac Myocyte Hypertrophy Signaling. J Biol Chem. 2012;287:42259–42268. doi:10.1074/jbc.M112.382937.

32. Zeigler AC, Richardson WJ, Holmes JW, Saucerman JJ. A Computational Model of Cardiac Fibroblast Signaling Predicts Context-Dependent Drivers of Myofibroblast Differentiation. J Mol Cell Cardiol. 2016;94:72–81. doi:10.1016/j.yjmcc.2016.03.008.

33. Zapotoczny B, Szafranska K, Owczarczyk K, Kus E, Chlopicki S, Szymonski M. Atomic Force Microscopy Reveals the Dynamic Morphology of Fenestrations in Live Liver Sinusoidal Endothelial Cells. Sci Rep. 2017;7:7994. doi:10.1038/s41598-017-08555-0.

34. Shu A, Du Q, Chen J, Gao Y, Zhu Y, Lv G, et al. Catalpol ameliorates endothelial dysfunction and inflammation in diabetic nephropathy via suppression of RAGE/RhoA/ROCK signaling pathway. Chem Biol Interact. 2021;348:109625. doi:10.1016/j.cbi.2021.109625.

35. Svistounov D, Warren A, McNerney GP, Owen DM, Zencak D, Zykova SN, et al. The Relationship between Fenestrations, Sieve Plates and Rafts in Liver Sinusoidal Endothelial Cells. PLoS ONE. 2012;7:e46134. doi:10.1371/journal.pone.0046134.

36. Mou X, Leeman SM, Roye Y, Miller C, Musah S. Fenestrated Endothelial Cells across Organs: Insights into Kidney Function and Disease. Int J Mol Sci. 2024;25:9107. doi:10.3390/ijms25169107.

37. Weil E, Lemley KV, Mason CC, Yee B, Jones LI, Blouch K, et al. Podocyte detachment and reduced glomerular capillary endothelial fenestration promote kidney disease in type 2 diabetic nephropathy. Kidney Int. 2012; p. 1010–1017. doi:10.1038/ki.2012.234.

38. Howsmon DP, Sacks MS. On Valve Interstitial Cell Signaling: The Link between Multiscale Mechanics and Mechanobiology. Cardiovasc Eng Tech. 2021;12:15–27. doi:10.1007/s13239-020-00509-4.

39. He FF, Chen S, Su H, Meng XF, Zhang C. Actin-Associated Proteins in the Pathogenesis of Podocyte Injury. Curr Genomics. 2013;14:477–484. doi:10.2174/13892029113146660014.

40. Blanchoin L, Boujemaa-Paterski R, Sykes C, Plastino J. Actin Dynamics, Architecture, and Mechanics in Cell Motility. Physiol Rev. 2014;94:235–263. doi:10.1152/physrev.00018.2013.

41. Mönkemöller V, Øie C, Hübner W, Huser T, McCourt P. Multimodal Super-Resolution Optical Microscopy Visualizes the Close Connection between Membrane and the Cytoskeleton in Liver Sinusoidal Endothelial Cell Fenestrations. Sci Rep. 2015;5:16279. doi:10.1038/srep16279.

42. Kraeutler MJ, Soltis AR, Saucerman JJ. Modeling cardiac β-adrenergic signaling with normalized-Hill differential equations: comparison with a biochemical model. BMC Syst Biol. 2010;4:157. doi:10.1186/1752-0509-4-157.

43. Clark AP, Chowkwale M, Paap A, Dang S, Saucerman JJ. Logic-Based Modeling of Biological Networks with Netflux. bioRxiv. 2024;Preprint:2024.01.11.575227. doi:10.1101/2024.01.11.575227.

44. Klinke II DJ, Finley SD. Timescale Analysis of Rule-Based Biochemical Reaction Networks. Biotechnol Prog. 2012;28:33–44. doi:10.1002/btpr.704.

45. Lee VK, Hosking BM, Holeniewska J, Kubala EC, Lundh von Leithner P, Gardner PJ, et al. BTBR Ob/Ob Mouse Model of Type 2 Diabetes Exhibits Early Loss of Retinal Function and Retinal Inflammation Followed by Late Vascular Changes. Diabetologia. 2018;61:2422–2432. doi:10.1007/s00125-018-4696-x.

46. Helton JC, Johnson JD, Sallaberry CJ, Storlie CB. Survey of sampling-based methods for uncertainty and sensitivity analysis. Reliab Eng Syst Saf. 2006;91:1175–1209. doi:10.1016/j.ress.2005.11.017.

47. Linden NJ, Kramer B, Rangamani P. Bayesian parameter estimation for dynamical models in systems biology. PLoS Comput Biol. 2022;18:1–48. doi:10.1371/journal.pcbi.1010651.

48. Hespanhol L, Vallio C, Saragiotto B, Costa L. Understanding and interpreting confidence and credible intervals around effect estimates. Braz J Phys Ther. 2018;23:290–301. doi:10.1016/j.bjpt.2018.12.006.

49. Huber HA, Georgia SK, Finley SD. Systematic Bayesian posterior analysis guided by Kullback-Leibler divergence facilitates hypothesis formation. J Theor Biol. 2023;558:111341. doi:10.1016/j.jtbi.2022.111341.

50. Student. The probable error of a mean. Biometrika. 1908; p. 1–25.

51. Stoehr JP, Byers JE, Clee SM, Lan H, Boronenkov IV, Schueler KL, et al. Identification of Major Quantitative Trait Loci Controlling Body Weight Variation in Ob/Ob Mice. Diabetes. 2004;53:245–249. doi:10.2337/diabetes.53.1.245.

52. Hudkins KL, Pichaiwong W, Wietecha T, Kowalewska J, Banas MC, Spencer MW, et al. BTBR Ob/Ob Mutant Mice Model Progressive Diabetic Nephropathy. J Am Soc Nephrol. 2010;21:1533–1542. doi:10.1681/ASN.2009121290.

53. Björnson Granqvist A, Ericsson A, Sanchez J, Tonelius P, William-Olsson L, Dahlqvist U, et al. High-Protein Diet Accelerates Diabetes and Kidney Disease in the BTBR ob/ob Mouse. Am J Physiol Renal Physiol. 2020;318:F763–F771. doi:10.1152/ajprenal.00484.2019.

54. Estrada AC, Yoshida K, Saucerman JJ, Holmes JW. A Multiscale Model of Cardiac Concentric Hypertrophy Incorporating Both Mechanical and Hormonal Drivers of Growth. Biomech Model Mechanobiol. 2021;20:293–307. doi:10.1007/s10237-020-01385-6.

55. Cassis P, Zoja C, Perico L, Remuzzi G. A preclinical overview of emerging therapeutic targets for glomerular diseases. Expert Opin Ther Targets. 2019;23:593–606. doi:10.1080/14728222.2019.1626827.

