## Supplementary material for "Multi-Cellular Network Model Predicts Alterations in Glomerular Endothelial Structure in Diabetic Kidney Disease": S1 Appendix

#### Supplementary equations

Besides the equations reported in the main text, the following LBODEs (Eqs A1–A35) also govern network interactions.

$$\frac{d\mathrm{AGE}}{dt} = \frac{y_{\max_{\mathrm{AGE}}} f_{\mathrm{act}_3}(\mathrm{GLU}) - \mathrm{AGE}}{\tau_{\mathrm{AGE}}} \tag{A1}$$

$$\frac{d\mathrm{VEGFR1}}{dt} = \frac{y_{\max_{\mathrm{VEGFR1}}} f_{\mathrm{act}_{18}}(\mathrm{VEGF\text{-}A}) - \mathrm{VEGFR1}}{\tau_{\mathrm{VEGFR1}}} \tag{A2}$$

$$\frac{d\mathrm{VEGFR2}}{dt} = \frac{y_{\max_{\mathrm{VEGFR2}}} f_{\mathrm{act}_{19}}(\mathrm{VEGF\text{-}A}) - \mathrm{VEGFR2}}{\tau_{\mathrm{VEGFR2}}} \tag{A3}$$

$$\frac{d\mathrm{VEGF\text{-}A_{mRNA}}}{dt} = \frac{y_{\max_{\mathrm{VEGF\text{-}A_{mRNA}}}} f_{\mathrm{act}_{15}}(\mathrm{NF\kappa B}) - \mathrm{VEGF\text{-}A_{mRNA}}}{\tau_{\mathrm{VEGF\text{-}A_{mRNA}}}} \tag{A4}$$

$$\frac{d\mathrm{RAGE}}{dt} = \frac{y_{\max_{\mathrm{RAGE}}} f_{\mathrm{act}_4}(\mathrm{AGE}) - \mathrm{RAGE}}{\tau_{\mathrm{RAGE}}} \tag{A5}$$

$$\frac{d\mathrm{RAGE_{ec}}}{dt} = \frac{y_{\max_{\mathrm{RAGE_{ec}}}} f_{\mathrm{act}_{20}}(\mathrm{AGE}) - \mathrm{RAGE_{ec}}}{\tau_{\mathrm{RAGE_{ec}}}} \tag{A6}$$

$$\frac{d\mathrm{IL\text{-}1R}}{dt} = \frac{y_{\max_{\mathrm{IL\text{-}1R}}} f_{\mathrm{act}_2}(\mathrm{IL\text{-}1\beta}) - \mathrm{IL\text{-}1R}}{\tau_{\mathrm{IL\text{-}1R}}} \tag{A7}$$

$$\frac{d\mathrm{NADPH}}{dt} = \frac{y_{\max_{\mathrm{NADPH}}} f_{\mathrm{act}_5}(\mathrm{RAGE}) - \mathrm{NADPH}}{\tau_{\mathrm{NADPH}}} \tag{A8}$$

$$\frac{d\mathrm{NADPH_{ec}}}{dt} = \frac{y_{\max_{\mathrm{NADPH_{ec}}}} f_{\mathrm{act}_{21}}(\mathrm{RAGE_{ec}}) - \mathrm{NADPH_{ec}}}{\tau_{\mathrm{NADPH_{ec}}}} \tag{A9}$$

$$\frac{d\mathrm{ROS}}{dt} = \frac{y_{\max_{\mathrm{ROS}}} OR\left(f_{\mathrm{act}_{11}}(\mathrm{PI3K}), f_{\mathrm{act}_8}(\mathrm{NADPH})\right) - \mathrm{ROS}}{\tau_{\mathrm{ROS}}} \tag{A10}$$

$$\frac{d\mathrm{ROS_{ec}}}{dt} = \frac{y_{\max_{\mathrm{ROS_{ec}}}} OR(f_{\mathrm{act}_{33}}(\mathrm{eNOS}), f_{\mathrm{act}_{25}}(\mathrm{NADPH_{ec}})) - \mathrm{ROS_{ec}}}{\tau_{\mathrm{ROS_{ec}}}} \tag{A11}$$

$$\frac{d\mathrm{PI3K}}{dt} = \frac{y_{\max_{\mathrm{PI3K}}} f_{\mathrm{act}_7}(\mathrm{IL} - 1\mathrm{R}) - \mathrm{PI3K}}{\tau_{\mathrm{PI3K}}} \tag{A12}$$

$$\frac{d\mathrm{AKT}}{dt} = \frac{y_{\max_{\mathrm{AKT}}} f_{\mathrm{act}_9}(\mathrm{PI3K}) - \mathrm{AKT}}{\tau_{\mathrm{AKT}}} \tag{A13}$$

$$\frac{d\mathrm{PI3K_{ec}}}{dt} = \frac{y_{\max_{\mathrm{PI3K_{ec}}}} OR(f_{\mathrm{act}_{23}}(\mathrm{VEGFR1}), f_{\mathrm{act}_{22}}(\mathrm{VEGFR2})) - \mathrm{PI3K_{ec}}}{\tau_{\mathrm{PI3K_{ec}}}} \tag{A14}$$

$$\frac{d\mathrm{AKT_{ec}}}{dt} = \frac{y_{\max_{\mathrm{AKT_{ec}}}} f_{\mathrm{act}_{25}}(\mathrm{PI3K_{ec}}) - \mathrm{AKT_{ec}}}{\tau_{\mathrm{AKT_{ec}}}} \tag{A15}$$

$$\frac{d\mathrm{NF\kappa B}}{dt} = \frac{y_{\max_{\mathrm{NF\kappa B}}} OR(AND(f_{\mathrm{act}_6}(\mathrm{IL\text{-}1R}), f_{\mathrm{act}_6}(\mathrm{ROS})), f_{\mathrm{act}_{12}}(\mathrm{AKT})) - \mathrm{NF\kappa B}}{\tau_{\mathrm{NF\kappa B}}} \tag{A16}$$

$$\frac{d\mathrm{NF\kappa B_{ec}}}{dt} = \frac{y_{\max_{\mathrm{NF\kappa B_{ec}}}} OR(f_{\mathrm{act}_{28}}(\mathrm{PLC\text{-}\gamma}), f_{\mathrm{act}_{29}}(\mathrm{ROS_{ec}})) - \mathrm{NF\kappa B_{ec}}}{\tau_{\mathrm{NF\kappa B_{ec}}}} \tag{A17}$$

$$\frac{d\mathrm{NO}}{dt} = \frac{y_{\max_{\mathrm{NO}}} OR(f_{\mathrm{act}_{32}}(\mathrm{eNOS}), f_{\mathrm{act}_{39}}(\mathrm{Ca})) - \mathrm{NO}}{\tau_{\mathrm{NO}}} \tag{A18}$$

$$\frac{d\text{ONOO}}{dt} = \frac{y_{\max_{\text{ONOO}}} AND(f_{\text{act}_{34}}(\text{NO}), f_{\text{act}_{34}}(\text{ROS}_{\text{ec}})) - \text{ONOO}}{\tau_{\text{ONOO}}} \tag{A19}$$

$$\frac{d\text{eNOS}}{dt} = \frac{y_{\max_{\text{eNOS}}} f_{\text{act}_{28}}(\text{AKT}_{\text{ec}}) - \text{eNOS}}{\tau_{\text{eNOS}}} \tag{A20}$$

$$\frac{d\text{IL-6}}{dt} = \frac{y_{\max_{\text{IL-6}}} OR(f_{\text{act}_{13}}(\text{NF}\kappa\text{B}), f_{\text{act}_{30}}(\text{NF}\kappa\text{B}_{\text{ec}})) - \text{IL-6}}{\tau_{\text{IL-6}}} \tag{A21}$$

$$\frac{d\text{TNF-}\alpha}{dt} = \frac{y_{\max_{\text{TNF-}\alpha}} OR(f_{\text{act}_{14}}(\text{NF}\kappa\text{B}), f_{\text{act}_{11}}(\text{NF}\kappa\text{B}_{\text{ec}})) - \text{TNF-}\alpha}{\tau_{\text{TNF-}\alpha}} \tag{A22}$$

$$\frac{d\text{IL-}\beta}{dt} = \frac{y_{\max_{\text{IL-1}\beta}} OR(f_{\text{act}_{17}}(\text{NF}\kappa\text{B}), f_{\text{act}_{31}}(\text{NF}\kappa\text{B}_{\text{ec}})) - \text{IL-1}\beta}{\tau_{\text{IL-1}\beta}} \tag{A23}$$

$$\frac{d\text{PLC-}\gamma}{dt} = \frac{y_{\max_{\text{PLC-}\gamma}} f_{\text{act}_{27}}(\text{VEGFR1}) - \text{PLC-}\gamma}{\tau_{\text{PLC-}\gamma}} \tag{A24}$$

$$\frac{d\text{VEGF-A}}{dt} = \frac{y_{\max_{\text{VEGF-A}}} f_{\text{act}_{16}}(\text{VEGF-A}_{\text{mRNA}}) - \text{VEGF-A}}{\tau_{\text{VEGF-A}}} \tag{A25}$$

$$\frac{d\text{Ca}}{dt} = \frac{y_{\max_{\text{Ca}}} OR(f_{\text{inhib}_{35}}(\text{NO}), f_{\text{act}_{36}}(\text{PLC-}\gamma)) - \text{Ca}}{\tau_{\text{Ca}}} \tag{A26}$$

$$\frac{d\text{pJunction}}{dt} = \frac{y_{\max_{\text{pJunction}}} f_{\text{act}_{37}}(\text{Ca}) - \text{pJunction}}{\tau_{\text{pJunction}}} \tag{A27}$$

$$\frac{d\text{Gap Width}}{dt} = \frac{y_{\max_{\text{Gap Width}}} f_{\text{act}_{38}}(\text{pJunction}) - \text{Gap Width}}{\tau_{\text{Gap Width}}} \tag{A28}$$

$$\frac{d\text{Actin}_{\text{r}}}{dt} = \frac{y_{\max_{\text{Actin}_{\text{r}}}} f_{\text{act}_{48}}(\text{MLC}) - \text{Actin}_{\text{r}}}{\tau_{\text{Actin}_{\text{r}}}} \tag{A29}$$

$$\frac{d\text{Actin}_{\text{s}}}{dt} = \frac{y_{\max_{\text{Actin}_{\text{s}}}} W_{\text{pMLC}} - \text{Actin}_{\text{s}}}{\tau_{\text{Actin}_{\text{s}}}} \tag{A30}$$

$$\frac{d\text{RhoRock}}{dt} = \frac{y_{\max_{\text{RhoRock}}} f_{\text{act}_{44}}(\text{VEGFR2}) - \text{RhoRock}}{\tau_{\text{RhoRock}}} \tag{A31}$$

$$\frac{d\text{MLCK}}{dt} = \frac{y_{\max_{\text{MLCK}}} OR(f_{\text{inhib}_{41}}(\text{Ca}), AND(f_{\text{inhib}_{40}}(\text{NO}), f_{\text{act}_{40}}(\text{ROS}))) - \text{MLCK}}{\tau_{\text{MLCK}}} \tag{A32}$$

$$\frac{d\text{pMLC}}{dt} = \frac{y_{\max_{\text{pMLC}}} OR(f_{\text{act}_{42}}(\text{RhoRock}), AND(f_{\text{act}_{46}}(\text{MLC}), f_{\text{act}_{46}}(\text{MLCK})) - \text{pMLC}}{\tau_{\text{pMLC}}} \tag{A33}$$

$$\frac{d\text{MLCP}}{dt} = \frac{y_{\max_{\text{MLCP}}} f_{\text{inhib}_{45}}(\text{RhoRock}) - \text{MLCP}}{\tau_{\text{MLCP}}} \tag{A34}$$

$$\frac{d\text{MLC}}{dt} = \frac{y_{\max_{\text{MLC}}} AND(f_{\text{act}_{43}}(\text{pMLC}), f_{\text{act}_{43}}(\text{MLCP})) - \text{MLC}}{\tau_{\text{MLC}}} \tag{A35}$$

#### Supplementary tables

**Table A1.** Chemical species abbreviations and definitions used in the extended network model.

| Abbreviation | Definition |
|---|---|
| AGE | advanced glycation end product |
| AKT | serine/threonine-specific protein kinases |
| Ca | calcium |
| eNOS | endothelial nitric oxide synthase |
| Gap Width | intercellular gap width between GECs |
| GLU | glucose |
| IL | interleukin |
| MLCK | myosin light chain kinase |
| MLCP | myosin light chain phosphotase |
| NADPH | nicotinamide adenine dinucleotide phosphate |
| NF$\kappa$B | nuclear factor kappa B |
| NO | nitric oxide |
| ONOO | peroxynitrite |
| PI3K | phosphoinositide 3-kinases |
| pJunction | phosphorylated junction protein |
| PLC-$\gamma$ | phospholipase C gamma |
| RAGE | receptor of advanced glycation end product |
| RhoRock | Rho-associated protein kinase |
| ROS | reactive oxygen species |
| TLR | toll-like receptor |
| TNF-$\alpha$ | tumor necrosis factor-alpha |
| VEGF | vascular endothelial growth factor |
| VEGFR | vascular endothelial growth factor receptor |

**Table A2.** List of species parameter $\tau_i$ for the LBODEs model organized by species index $i$. The time constant ($\tau_i$) values in units of hours are set based on realistic timescales for activation of respective protein [1]. The default value of $\tau_i$ is 1 hour. The default values of the other species parameters are initial value $y_{0_i} = 0$ and maximal value $y_{\max_i} = 1$ for all species unless stated otherwise. Number and Diameter represent fenestration number and fenestration diameter, respectively.

| ID | Species | $\tau_i$ (hour) | $y_{\max_i}$ | $y_{0_i}$ |
|---|---|---|---|---|
| 1 | GLU | 1 | 1 | 0 |
| 2 | $\text{Actin}_\text{s}$ | 1 | 1 | 0 |
| 3 | AGE | 1 | 1 | 0 |
| 4 | VEGFR1 | 0.35 | 1 | 0 |
| 5 | VEGFR2 | 0.35 | 1 | 0 |
| 6 | $\text{VEGF-A}_\text{mRNA}$ | 88 | 1 | 0 |
| 7 | $\text{RAGE}_\text{ec}$ | 0.35 | 1 | 0 |
| 8 | RAGE | 0.35 | 1 | 0 |
| 9 | IL-1R | 0.35 | 1 | 0 |
| 10 | NADPH | 1 | 1 | 0 |
| 11 | $\text{NADPH}_\text{ec}$ | 1 | 1 | 0 |
| 12 | $\text{ROS}_\text{ec}$ | 1 | 1 | 0 |
| 13 | ROS | 1 | 1 | 0 |
| 14 | PI3K | 1 | 1 | 0 |
| 15 | AKT | 1 | 1 | 0 |
| 16 | $\text{PI3K}_\text{ec}$ | 1 | 1 | 0 |
| 17 | $\text{AKT}_\text{ec}$ | 1 | 1 | 0 |
| 18 | $\text{NF}\kappa\text{B}_\text{ec}$ | 0.055 | 1 | 0 |
| 19 | NFκB | 0.055 | 1 | 0 |
| 20 | NO | 1 | 1 | 0 |
| 21 | ONOO | 1 | 1 | 0 |
| 22 | eNOS | 1 | 1 | 0 |
| 23 | IL-6 | 90 | 1 | 0 |
| 24 | TNF-α | 90 | 1 | 0 |
| 25 | IL-1β | 90 | 1 | 0 |
| 26 | PLC-γ | 1 | 1 | 0 |
| 27 | VEGF-A | 1.13 | 1 | 0 |
| 28 | pJunction | 1 | 1 | 0 |
| 29 | Ca | 1 | 1 | 0 |
| 30 | Gap Width | 1 | 1 | 0 |
| 31 | $\text{Actin}_\text{r}$ | 1 | 1 | 0 |
| 32 | RhoRock | 1 | 1 | 0 |
| 33 | MLCK | 1 | 1 | 0 |
| 34 | pMLC | 400.5 | 1 | 0 |
| 35 | MLCP | 1 | 1 | 0 |
| 36 | MLC | 1 | 1 | 0 |
| 37 | Number | – | – | 6.3 |
| 38 | Diameter | – | – | 47.9 nm |

**Table A3.** List of optimal reaction parameters for the LBODEs model organized by reaction index $j$. Reaction weight ($W_j$), Hill coefficient ($n_j$), and half-maximal effect ($EC_{50_j}$) are shown for respective reaction rules or interactions. Reaction rule A ⇒ C denotes input A activates output C. Reaction rule !A ⇒ C denotes input A inhibits output C. Reaction rule A & B ⇒ C denotes the $AND$ logic operator between input A and input B to give output C. Default values are $W_j = 1$, $n_j = 1.4$, and $EC_{50_j} = 0.5$. (–) indicates that the respective parameters are not applicable to the reaction.

| ID | Reaction | $W_j$ | $n_j$ | $EC_{50_j}$ |
|---|---|---|---|---|
| 1 | ⇒ GLU | – | – | – |
| 2 | IL-1β ⇒ IL-1R | 1 | 1.4 | 0.5 |
| 3 | GLU ⇒ AGE | 1 | 1.45 | 0.5 |
| 4 | AGE ⇒ RAGE | 1 | 2.71 | 0.474 |
| 5 | RAGE ⇒ NADPH | 0.944 | 2.7 | 0.470 |
| 6 | IL-1R & ROS ⇒ NFκB | 1 | 1.4 | 0.5 |
| 7 | IL-1R ⇒ PI3K | 1 | 1.4 | 0.5 |
| 8 | NADPH ⇒ ROS | 0.943 | 2.64 | 0.545 |
| 9 | PI3K ⇒ AKT | 1 | 2.7 | 0.5 |
| 10 | PI3K ⇒ ROS | 0.950 | 2.7 | 0.419 |
| 11 | $NF\kappa B_{ec}$ ⇒ TNF-α | 0.999 | 3.96 | 0.840 |
| 12 | AKT ⇒ NFκB | 1 | 2.7 | 0.5 |
| 13 | NFκB ⇒ IL-6 | 0.950 | 2.7 | 0.420 |
| 14 | NFκB ⇒ TNF-α | 0.950 | 2.7 | 0.419 |
| 15 | NFκB ⇒ $VEGF\text{-}A_{mRNA}$ | 0.95 | 2.7 | 0.422 |
| 16 | $VEGF\text{-}A_{mRNA}$ ⇒ VEGF-A | 0.949 | 2.77 | 0.595 |
| 17 | NFκB ⇒ IL-1β | 0.950 | 2.7 | 0.421 |
| 18 | VEGF-A ⇒ VEGFR1 | 1 | 2.71 | 0.5 |
| 19 | VEGF-A ⇒ VEGFR2 | 1 | 2.72 | 0.5 |
| 20 | AGE ⇒ RAGE | 1 | 1.56 | 0.839 |
| 21 | $RAGE_{ec}$ ⇒ $NADPH_{ec}$ | 0.938 | 1.6 | 0.839 |
| 22 | VEGFR2 ⇒ $PI3K_{ec}$ | 1 | 2.73 | 0.5 |
| 23 | VEGFR1 ⇒ $PI3K_{ec}$ | 1 | 2.72 | 0.5 |
| 24 | $NADPH_{ec}$ ⇒ $ROS_{ec}$ | 0.946 | 3.91 | 0.838 |
| 25 | $PI3K_{ec}$ ⇒ $AKT_{ec}$ | 1 | 2.93 | 0.688 |
| 26 | $AKT_{ec}$ ⇒ eNOS | 1 | 3.11 | 0.576 |
| 27 | VEGFR1 ⇒ PLC | 1 | 1.4 | 0.5 |
| 28 | PLC ⇒ $NFKB_{ec}$ | 1 | 1.4 | 0.5 |
| 29 | $ROS_{ec}$ ⇒ $NFKB_{ec}$ | 1 | 2.7 | 0.010 |
| 30 | $NFKB_{ec}$ ⇒ IL-6 | 0.986 | 2.7 | 0.471 |
| 31 | $NFKB_{ec}$ ⇒ IL-1β | 0.962 | 2.7 | 0.391 |
| 32 | eNOS ⇒ NO | 1 | 1.46 | 0.157 |
| 33 | eNOS ⇒ $ROS_{ec}$ | 1 | 3.66 | 0.833 |
| 34 | $ROS_{ec}$ & NO ⇒ ONOO | 1 | 1.4 | 0.5 |
| 35 | !NO ⇒ Ca | 1 | 2.68 | 0.5 |
| 36 | PLC-γ ⇒ Ca | 1 | 1.4 | 0.5 |
| 37 | Ca ⇒ pJunction | 1 | 1.4 | 0.5 |
| 38 | pJunction ⇒ Gap Width | 1 | 1.4 | 0.5 |
| 39 | Ca ⇒ NO | 1 | 1.4 | 0.5 |
| 40 | !NO & ROS ⇒ MLCK | 1 | 1.4 | 0.5 |
| 41 | Ca ⇒ MLCK | 1 | 1.4 | 0.5 |
| 42 | RhoRock ⇒ pMLC | 1 | 1.4 | 0.5 |
| 43 | pMLC & MLCP ⇒ MLC | 1 | 1.4 | 0.5 |
| 44 | VEGFR2 ⇒ RhoRock | 1 | 1.4 | 0.5 |
| 45 | !RhoRock ⇒ MLCP | 1 | 1.4 | 0.5 |
| 46 | MLC & MLCK ⇒ pMLC | 1 | 1.4 | 0.5 |
| 47 | pMLC ⇒ $Actin_s$ | 1 | 1.4 | 0.5 |
| 48 | MLC ⇒ $Actin_r$ | 1 | 1.4 | 0.5 |

**Table A4.** Experimentally observed effect (promotion ↑ or inhibition ↓) of chemical agents on respective targeted species in liver sinusoidal endothelial cells (LSEC) [2]. LSEC porosity is the number of fenestrations per unit area. LSEC diameter is the fenestration diameter.

| Chemical agent | Target | Effect on LSEC porosity | Effect on LSEC diameter |
|---|---|---|---|
| KN93 | Calcium | ↓ | no change |
| ML7 | MLCK | ↓ | ↑ |
| Y27632 | ROCK | ↑ | ↓ |
| Calyculin A | MLCP | ↓ | ↑ |
| Cytochalasin B | Stressed actin fibers | ↑ | ↑ |

### Supplementary figures

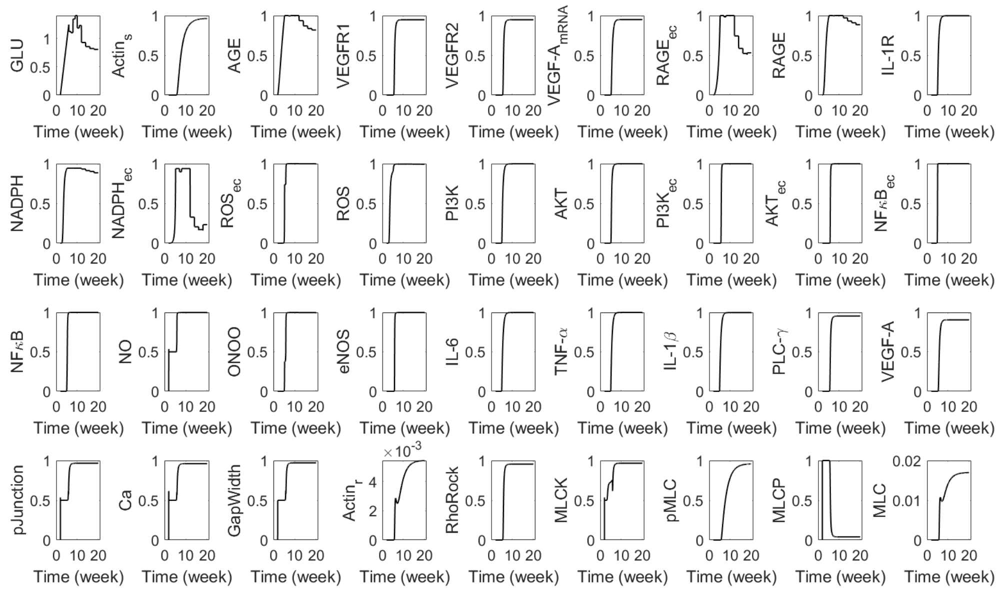


**Fig A1.** Predicted dynamics of species in the network simulated at glucose levels at the mean of observed data in diabetic mice (Fig 2).

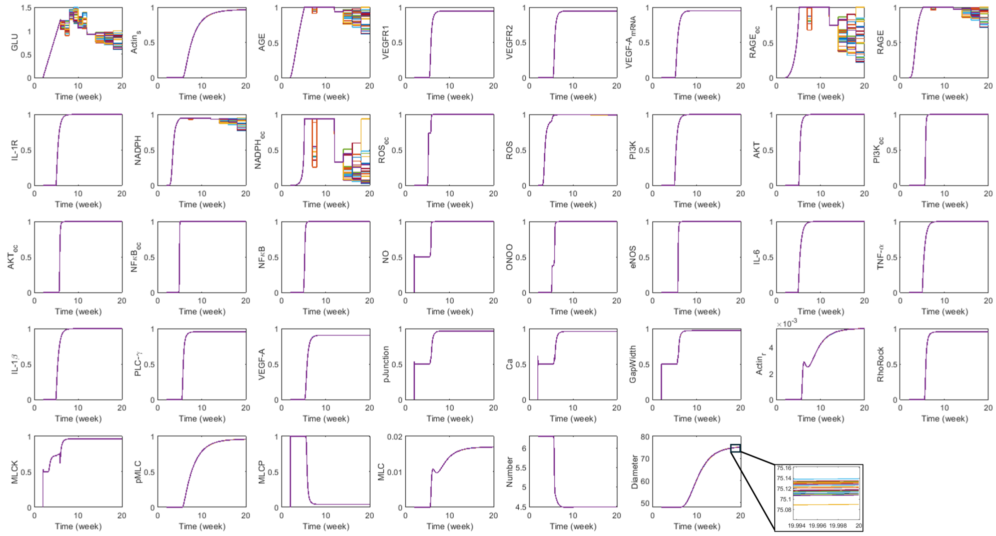

**Fig A2.** Predicted dynamics of species in the network for variable samples of glucose concentration within the error bars of observed data in diabetic mice.

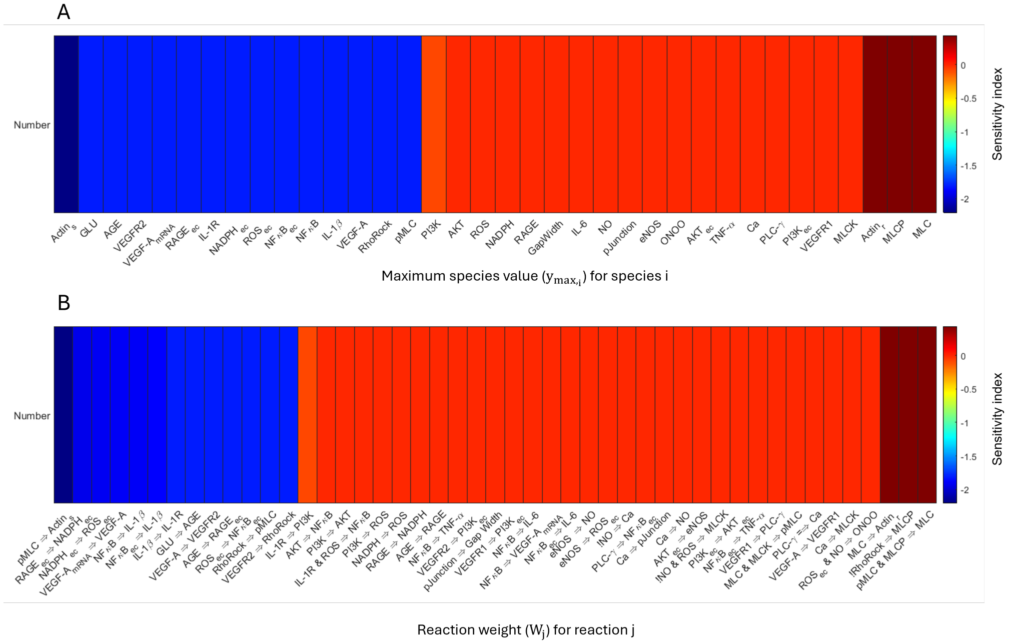

**Fig A3.** Calculated sensitivity indices (Eq 6) for fenestration number upon local perturbation in $y_{max_i}$ for respective species (horizontal axis, top panel) and upon local perturbation in $W_j$ for respective reaction rules (horizontal axis, bottom panel).

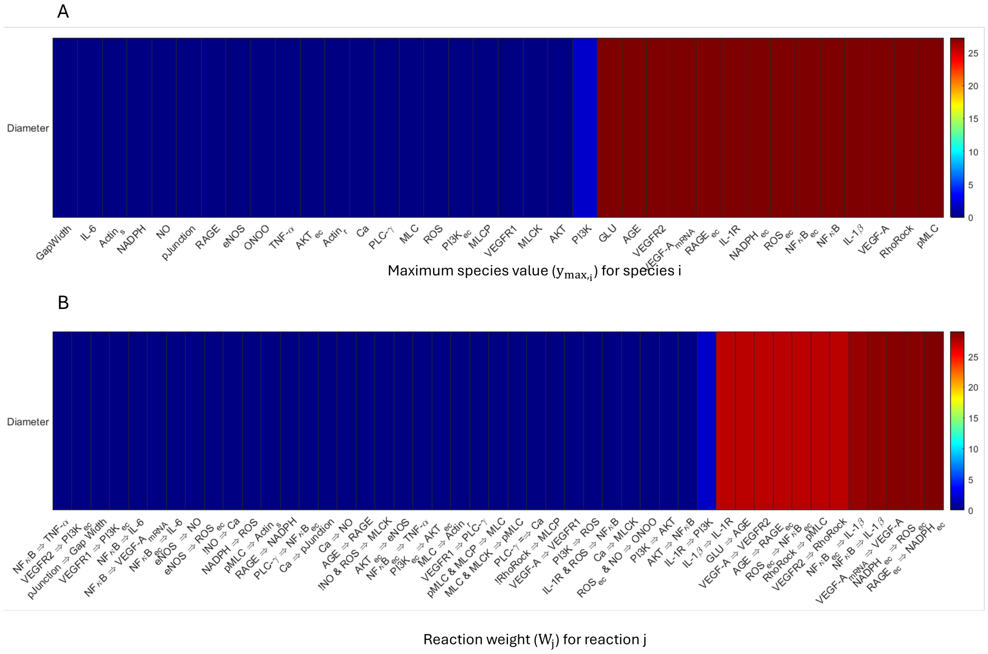


**Fig A4.** Calculated sensitivity indices (Eq 6) for fenestration diameter upon local perturbation in $y_{max_i}$ for respective species (horizontal axis, top panel) and upon local perturbation in $W_j$ for respective reaction rules (horizontal axis, bottom panel).

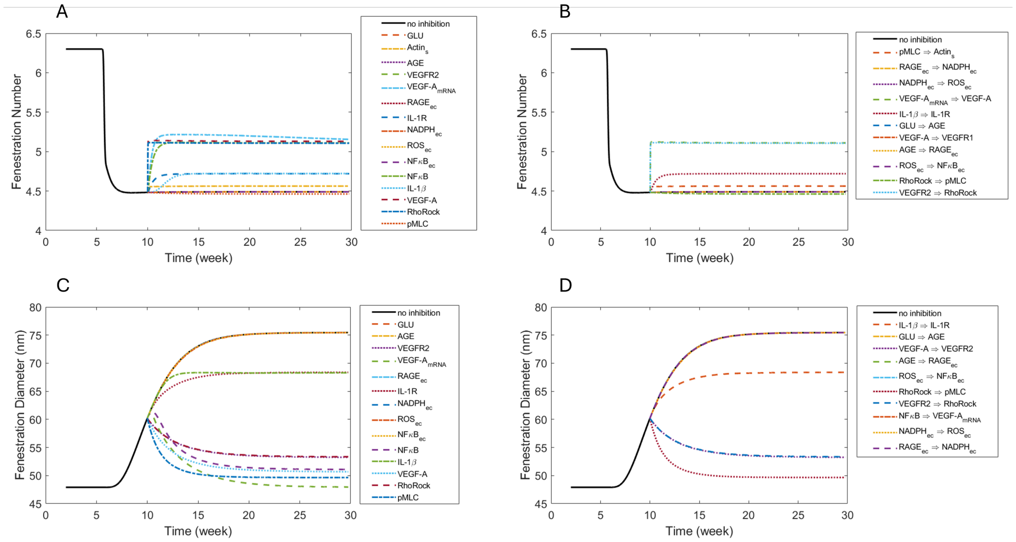


**Fig A5.** Effects at 10 weeks of 50% inhibition of sensitive (A) species and (B) reactions on fenestration number and of sensitive species (C) and (D) reactions on fenestration diameter.

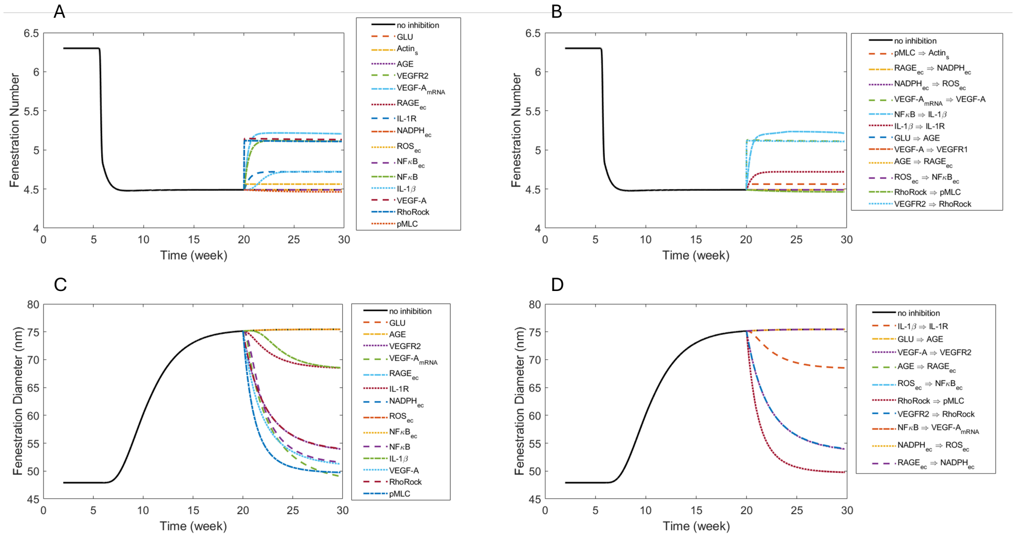


**Fig A6.** Effects at 20 weeks of 50% inhibition of sensitive (A) species and (B) reactions on fenestration number and of sensitive species (C) and (D) reactions on fenestration diameter.

### References


1. Klinke II DJ, Finley SD. Timescale Analysis of Rule-Based Biochemical Reaction Networks. Biotechnol Prog. 2012;28:33–44. doi:10.1002/btpr.704.
2. Zapotoczny B, Szafranska K, Lekka M, Ahluwalia BS, McCourt P. Tuning of liver sieve: the interplay between actin and myosin regulatory light chain regulates fenestration size and number in murine liver sinusoidal endothelial cells. Int J Mol Sci. 2022;23:9850. doi:10.3390/ijms23179850.
